# CRISPR-HAWK: Haplotype- and Variant-Aware Guide Design Toolkit for CRISPR-Cas

**DOI:** 10.64898/2025.12.27.696698

**Authors:** Alisa Kumbara, Manuel Tognon, Giulia Carone, Arianna Fontanesi, Nicola Bombieri, Rosalba Giugno, Luca Pinello

## Abstract

**Motivation:** Current CRISPR guide RNA design tools rely on reference genomes, overlooking how genetic variation impacts editing outcomes. As genome editing advances toward clinical applications, incorporating population diversity becomes essential for ensuring therapeutic efficacy across diverse populations.

**Results:** We present CRISPR-HAWK, a framework integrating individual- and population-scale variants and haplotypes into gRNA design. Analyzing therapeutic targets across 79,648 genomes reveals that genetic variants substantially alter guide performance. For the clinically approved sickle cell disease therapeutic guide targeting BCL11A, we identify haplotypes that completely abolish predicted cutting activity. Across seven therapeutic loci, 82.5% of guides contain variants modifying on-target activity. Variants also create novel protospacer adjacent motif sites generating individual-specific guides invisible to reference-based design. These findings demonstrate that variant-aware selection is critical for equitable genome editing.

**Availability:** CRISPR-HAWK is available at https://github.com/pinellolab/CRISPR-HAWK and https://github.com/InfOmics/CRISPR-HAWK

## 1 Introduction

CRISPR-Cas systems present unprecedented opportunities for therapeutic developments by offering a powerful means to precisely modify DNA within living cells. Originally repurposed from Streptococcus pyogenes Cas9 (SpCas9) for targeted genome editing, CRISPR has since evolved into a versatile platform comprising a diverse array of Cas orthologs and engineered variants (Jiang and Doudna, 2017). Central to this technology is a guide RNA (gRNA) that directs the Cas complex to a complementary genomic sequence, contingent on the presence of a protospacer adjacent motif (PAM). This modular mechanism enables a spectrum of biological outcomes, including the introduction of double-strand breaks, precise nucleotide substitution via base editors (Rees and Liu, 2018), combinations of genetic modifications via prime editing (Anzalone et al., 2019), and epigenetic or transcriptional modulation through CRISPRa/I systems (Thakore et al., 2016; Kampmann, 2018).

The success of CRISPR genome editing relies on two key factors: achieving high on-target efficiency while minimizing off-target activity (Clement et al., 2020). On-target efficiency refers to the ability of a gRNA to precisely direct the Cas nuclease to the intended genomic locus, thereby executing the desired genetic modification. Conversely, off-target effects occur when the CRISPR system binds and edits unintended genomic loci with partial sequence similarity, potentially resulting in harmful or confounding mutations (Cho et al., 2014). Several computational tools have been developed to support gRNA design by optimizing both aspects (Hanna and Doench, 2020). Tools such as Cas-Designer (Cho et al., 2014) or CHOPCHOP (Labun et al., 2019) identify candidate guides by aligning input PAM sequences against a reference genome. For each reported gRNA, they compute different sequence-based features, such as GC content, and estimate the number and location of potential off-target sites. This information is then used to rank guides based on predicted editing efficiency and target specificity. CRISPick (Doench et al., 2016; DeWeirdt et al., 2021) and CRISPRon (Anthon et al., 2022) integrate machine learning models trained on experimental data to prioritize guides with high predicted activity and minimal off-target potential. CRISPOR (Concordet and Haeussler, 2018), meanwhile, integrates information on single nucleotide polymorphisms (SNPs) that may overlap with candidate gRNA sequences.

However, existing tools have a critical limitation: gRNA selection is based on the reference genome, thereby potentially overlooking on-target sites introduced or modulated by genetic variants. While CRISPOR annotates candidate gRNAs with overlapping SNPs, the guide selection process still relies on the reference sequence. Consequently, the combined effect of multiple variants or haplotypes is not considered, nor is the impact of genetic variants on on-target activity quantitatively assessed. While it is well recognized that variants can create or modulate off-target sites (Cancellieri et al., 2023; Lazzarotto et al., 2025), there is growing evidence indicating that variants can affect on-target efficiency. This occurs through direct alteration of the intended protospacer sequences or changes in its local genomic context (Canver et al., 2018). Such effects are critical in therapeutic applications, where even subtle differences in editing efficiency or specificity can impact clinical efficacy and safety (Lessard et al., 2017; Liu et al., 2021).

The clinical significance of this problem is exemplified by recent therapeutic applications. Casgevy (exagamglogene autotemcel), one of the first FDA-approved CRISPR therapies for sickle cell disease and beta-thalassemia, relies on precise editing of the BCL11A enhancer using guide sg1617 (Frangoul et al., 2021). Yet these treatments, designed using reference genomes, may have variable efficacy across genetically diverse populations.

As genome editing advances toward personalized and clinical applications, the need to account for both individual- and population-level genetic variation becomes increasingly important (Scott and Zhang, 2017). Incorporating genetic diversity into gRNA design is key to enhancing on-target efficiency, reducing off-target effects, and ensuring the safety and efficacy of therapeutical outcomes.

To address these challenges, we developed CRISPR-HAWK, representing, to our knowledge, the first framework to integrate both genetic variants and haplotypes in gRNA design. By reconstructing individual haplotypes from genetic population-scale variant datasets, CRISPR-HAWK enables sample-specific guide selection while providing comprehensive assessment of guide performance. On-target efficiency of candidate gRNAs is predicted using machine learning models, while off-target activity is assessed using CRISPRitz (Cancellieri et al., 2020), a robust off-target nomination search engine. Candidate guides are further annotated with genetic variants, and functional information to support informed gRNA selection. We demonstrate the utility of CRISPR-HAWK, by designing candidate gRNAs targeting clinically relevant or widely tested genomic regions in human, while accounting for genetic variants from the 1000 Genomes Project (1000G) (Consortium et al., 2015; Zheng-Bradley et al., 2017), Human Genome Diversity Project (HGDP) (Bergström et al., 2020), and Genome Aggregation Database (gnomAD) (Karczewski et al., 2020; Chen et al., 2024) datasets. We show that genetic variation gives rise to numerous population- and individual-specific alternative gRNAs within each analyzed target region, revealing that many candidate guides are overlooked when relying solely on the reference genome. Moreover, we analyze how genetic variants impact on-target activity of gRNAs designed on the reference genome. These findings underscore the importance of adopting variant-aware strategies to ensure accurate, efficient, and equitable genome editing, particularly in therapeutic contexts.

## 2 Materials & Methods

CRISPR-HAWK is a command-line tool designed for the efficient enumeration, scoring, and annotation of CRISPR-Cas guide RNAs within user-defined genomic regions. **Figures 1** and **2** provide an overview of its architecture, illustrating the required and optional inputs, the main computational steps, and the structure of the resulting outputs. The following subsections detail the implementation of CRISPR-HAWK, including its strategy for integrating genetic variants and haplotype data, the algorithmic details underlying gRNA search and scoring, and the annotation pipeline. We further describe the datasets and experimental settings used for benchmarking and performance evaluation.

**Fig. 1:**
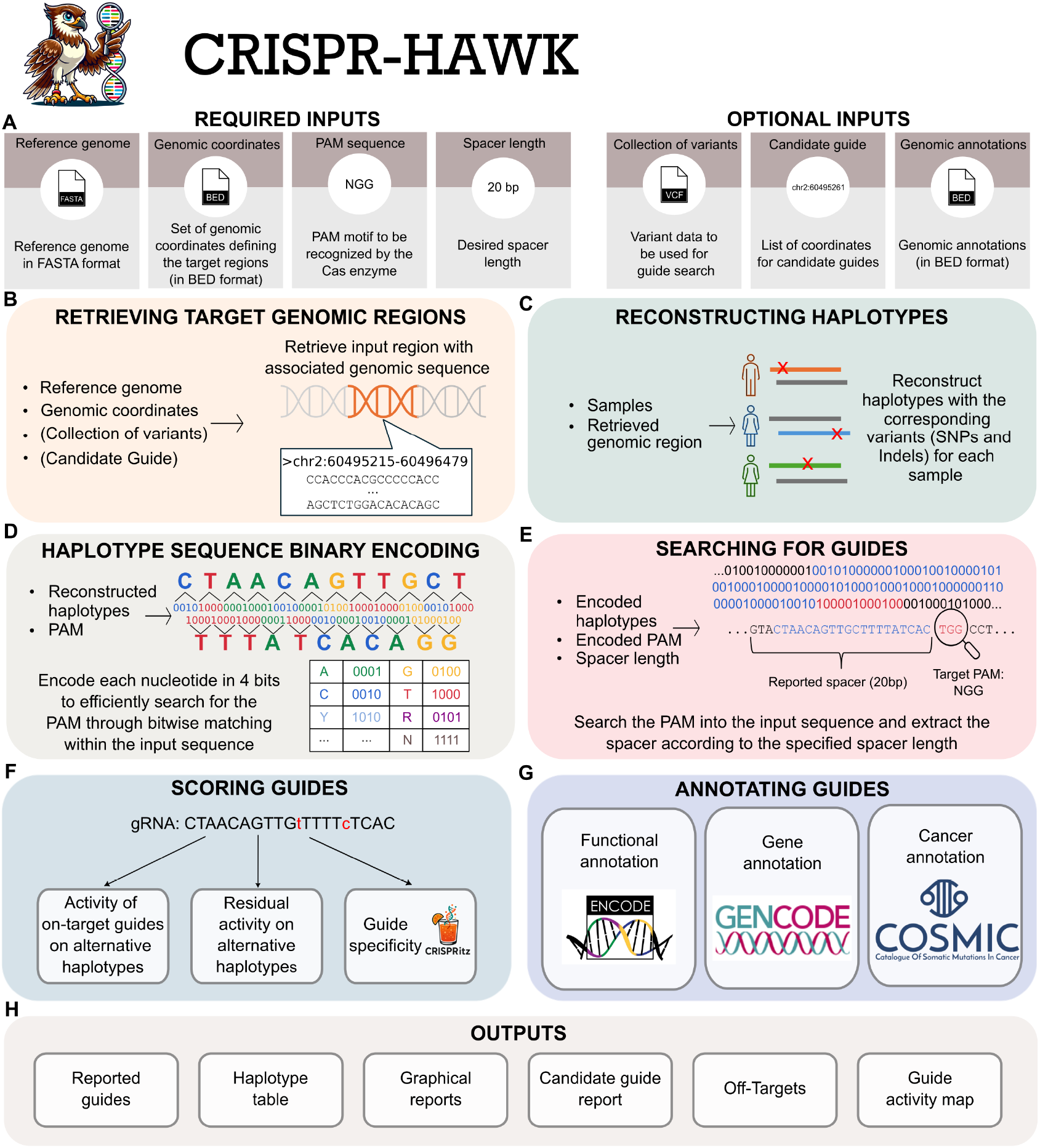
Overview of CRISPR-HAWK architecture. **(A)** CRISPR-HAWK designs gRNAs from a reference genome, by specifying genomic coordinates, a PAM motif, and a spacer length. Optional inputs include variant datasets for variant- and haplotype-aware guide design, genomic annotation files, and user-provided candidate guide coordinates for assessing variant effects. **(B)** Genomic regions of interest are extracted and **(C)** combined with variant data, when available, to reconstruct haplotype-resolved sequences. **(D)** Reconstructed haplotypes and PAM sequences are encoded using a 4-bit binary scheme that supports IUPAC ambiguity codes, enabling fast bitwise pattern matching. **(E)** Encoded sequences are scanned to identify PAM occurrences, and adjacent spacers are extracted to generate candidate gRNAs. (**F**) Candidate guides are evaluated using three complementary metrics: on-target efficiency of haplotype-matched guides, residual on-target activity of reference-designed guides on variant-containing target sequences, and guide specificity via existing predictive models. (**G**) Guides are annotated with functional, gene-level, and cancer-related features using default annotation tracks (ENCODE, GENCODE, COSMIC) or user-provided BED files. (**H**) CRISPR-HAWK generates comprehensive outputs including a table of all candidate guides identified within the input region. Optional outputs include a table of reconstructed haplotypes, graphical reports, a dedicated report for user-selected candidate guides, a genome-wide list of predicted off-target sites, and a guide activity map illustrating residual on-target activity with specificity.

**Fig. 2:**
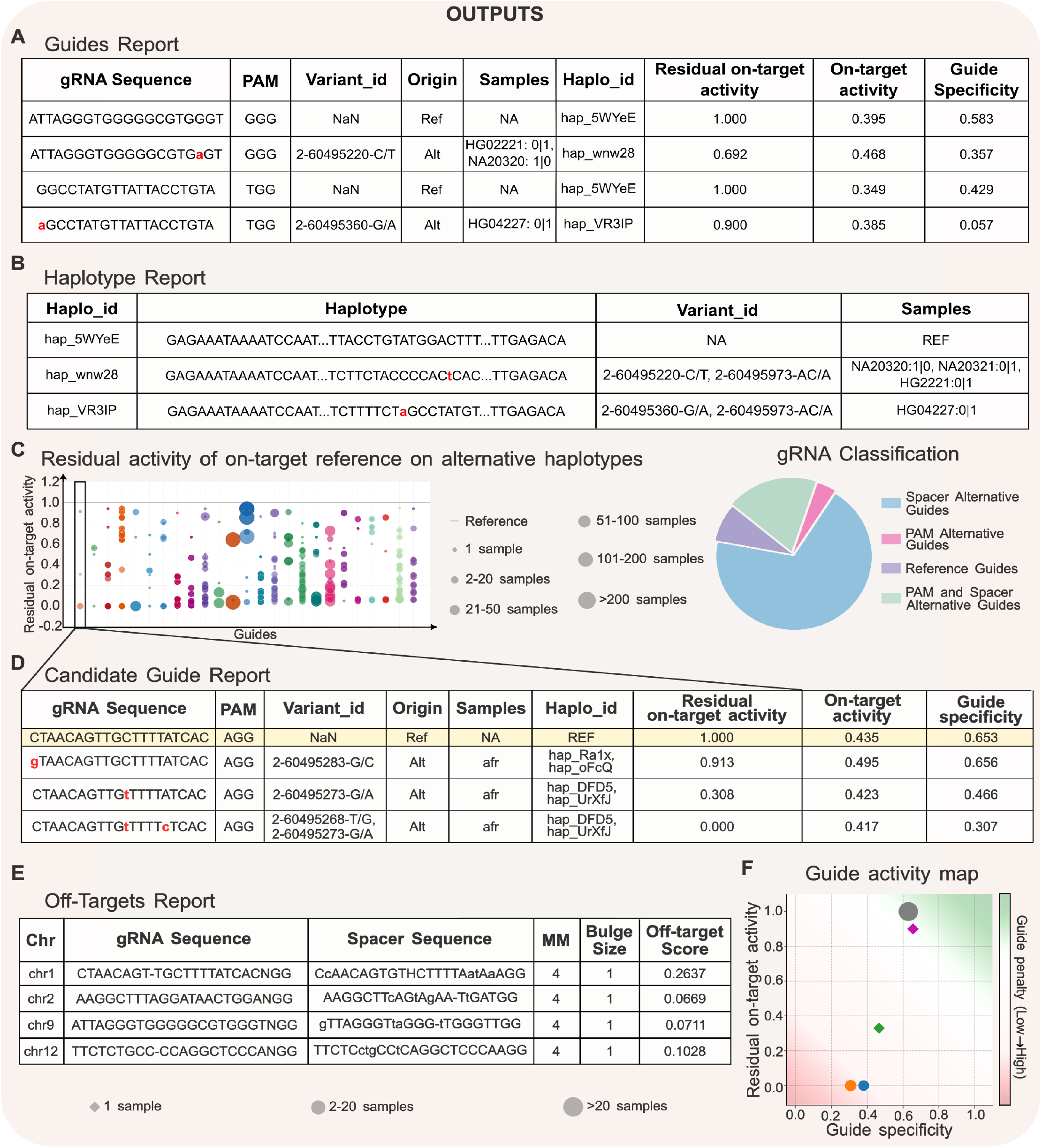
Overview of CRISPR-HAWK outputs and visualizations. **(A)** The primary output summarizes all candidate gRNAs with their associated performance metrics in a tabular report, including on-target efficiency, residual on-target activity, and guide specificity. **(B)** When requested, a haplotype table is provided, listing each reconstructed haplotype alongside the corresponding samples and variants. **(C)** Graphical reports include scatter plots depicting residual on-target activity across variant-containing target sequences, with point size indicating the number of individuals carrying each variant, and pie charts classifying gRNAs by type (reference, spacer alternative, PAM alternative, or both). **(D)** Users may optionally restrict the analysis to selected candidate guides. In this mode, the tool produces a dedicated report detailing all variant-containing target sequences and their associated scores for on-target efficiency, residual on-target activity, and guide specificity. **(E)** A genome-wide list of predicted off-target sites is generated for each candidate gRNA using CRISPRitz. **(F)** The guide activity map depicts residual on-target activity with guide specificity, providing a global view of guide performance across diverse genomic backgrounds.

### 2.1 Required and Optional Input Parameters

CRISPR-HAWK operates through a set of main input parameters guiding its search and annotation workflow (**Figure 1A**). Four inputs are mandatory: (i) a reference genome in FASTA format, (ii) genomic coordinates specifying the target regions in BED format, (iii) the PAM recognized by the selected Cas nuclease, and (iv) the desired spacer length. In addition to these core inputs, users may optionally provide variant datasets in VCF format (phased or unphased) to enable variant- and haplotype-aware guide design, as well as pre-defined candidate gRNAs for focused evaluation. Functional genomic annotation in BED format can also be provided to enrich the resulting guides with contextual information such as regulatory or coding region overlap.

### 2.2 Retrieving Target Genomic Regions and Haplotypes Reconstruction

Given a reference genome and a set of genomic coordinates, CRISPR-HAWK begins by extracting the corresponding target genomic regions from the reference genome (**Figure 1B**). To ensure sufficient context for both gRNA identification and downstream analyses, such as on-target efficiency prediction and variant mapping, each region is symmetrically extended by 100 base pairs upstream and downstream. The resulting extended sequences form the basis for integrating genetic variation and reconstructing haplotype-resolved representations of the target loci (**Figure 1C**). To incorporate genetic diversity, CRISPR-HAWK enriches the extracted sequences with single-nucleotide variants (SNVs) and short insertions/deletions (indels) provided in input VCF files. The tool supports both phased and unphased genotype data. For phased data, CRISPR-HAWK reconstructs haplotypes by creating two separate sequences per individual corresponding to the maternal and paternal alleles and sequentially inserts the appropriate variants. This process yields accurate reconstruction of each individual’s diploid genomic context within the target region. In the case of unphased data, where the chromosomal origin of each allele is unknown, heterozygous and multiallelic positions are initially encoded using IUPAC ambiguity codes which represent sets of possible nucleotides at each site. To resolve these ambiguities, CRISPR-HAWK performs combinatorial enumeration of all feasible haplotypes that could arise from the observed genotypes, only for sequences identified as guide candidates (see **Section 2.3**). This selective strategy significantly reduces memory consumption and computational time, enabling the analysis of large population-scale datasets, such as gnomAD. To further reduce computational burdens, CRISPR-HAWK collapses biologically equivalent haplotypes using hashing-based deduplication. Importantly, the haplotype reconstruction process considers the combined effect of SNVs and indels in both phased and unphased contexts, representing each individual’s sequence consistently. Each reconstructed haplotype is annotated with rich metadata including the list of occurring variants, samples or populations of origin, and phasing status (in case of phased variants only). This haplotype-centric approach enables CRISPR-HAWK to identify gRNAs with improved population coverage and individual-level precision.

### 2.3 Genomic Regions Binary Encoding and Search for Candidate Guides

Guide RNAs are identified by scanning the reconstructed haplotype sequences for the user-specified PAM motif on both forward and reverse strands. This step is computationally intensive, as it requires exhaustive base-by-base comparisons across potentially thousands of haplotypes. To address this, CRISPR-HAWK employs an optimized binary encoding strategy for guide discovery (**Figure 1D**). Each haplotype is converted into a compact 4-bits-per-base representation that supports both standard and IUPAC ambiguity codes. PAM detection is then performed via bitwise AND operations rather than character-level string comparisons. This approach reduces the constant-factor cost of the per-position matching step while decreasing the memory footprint required to store haplotype sequences relative to standard ASCII representation. Both the binary-encoded implementation and a naïve character-based scan share the same asymptotic time complexity *O*(*H* · *L*), where *H* denotes the number of haplotypes and *L* their length. The advantage of the binary representation therefore lies in improved constant-factor runtime and reduced memory footprint, which becomes increasingly important when analyzing large genomic regions or genetically diverse populations (see **Supplementary Data Section 1**). Upon detection of a valid PAM sequence, the adjacent spacer sequence, of user-defined length, is extracted (**Figure 1E**). For hits on the reverse strand the corresponding sequences are reverse-complemented to preserve consistent orientation. These extracted sequences are hereafter referred to as candidate gRNAs. To capture the impact of genetic variation, CRISPR-HAWK systematically flags all candidate gRNAs that overlap with variant sites. Within each guide, variant-affected nucleotides are displayed in lowercase, while reference-matching bases remain in uppercase, enabling clear distinction between conserved and variable positions. This representation facilitates rapid visual inspection and supports flexible downstream filtering, allowing users to prioritize guides based on their tolerance or sensitivity to genetic variation, depending on the intended application.

### 2.4 Scoring and Annotating Guides

Accurate evaluation of the designed gRNAs requires assessing their cleavage efficiency, specificity, and functional genomic context. CRISPR-HAWK leverages multiple scoring models (Figure 1F) and a comprehensive annotation framework (Figure 1G) to support informed guide selection.

We distinguish three complementary performance metrics. On-target efficiency predicts the cleavage activity of a guide at its intended target site, estimated using sequence-based machine learning models. Residual on-target activity predicts how effectively a reference-designed guide will cleave variant-containing target sequences, quantifying the impact of genetic mismatches on expected performance. Guide specificity measures selectivity across the genome by aggregating predicted off-target cleavage likelihoods. The following subsections describe each component in detail.

#### 2.4.1 Predicting On-Target Efficiency

Several computational models have been developed to predict on-target cleavage efficiency of gRNAs (Doench et al., 2014; Sherkatghanad et al., 2023). CRISPR-HAWK includes a diverse set of state-of-the-art models to provide robust and complementary estimates of guide activity.

For SpCas9, CRISPR-HAWK incorporates five scoring methods: Azimuth (Doench et al., 2016), Rule Set 3 (RS3) (DeWeirdt et al., 2022), CRISPRon (Anthon et al., 2022), sgDesigner (Hiranniramol et al., 2020), and PLM-CRISPR (Hou et al., 2025). Azimuth employs a machine learning model trained on more than 4,000 experimentally validated gRNAs targeting coding regions across 17 genes. RS3 extends Azimuth by incorporating additional sequence and thermodynamics features in the model, including poly(T) content, spacer-DNA melting temperature, and minimum free energy of the folded gRNA structure. CRISPRon is a deep learning model trained on over 23,000 gRNAs, combining sequence one-hot encoding with thermodynamic features to estimate on-target activity and prioritize candidate guides. sgDesigner, instead, was trained on a custom plasmid library comprising more than 7,400 sgRNAs. It employs a stacked generalization strategy integrating support vector machine and XGBoost models through logistic regression, leveraging both sequence and structural features. PLM-CRISPR is a dual-input deep learning framework that models sgRNA sequences efficiency on different Cas9 protein variants. While predicting gRNA activity, it accounts for sequence-protein interactions.

In addition, to support Cas12a applications, CRISPR-HAWK includes DeepCpf1 (Kim et al., 2018), a deep learning model trained to estimate the on-target activity of Cpf1 (Cas12a) systems using high-throughput experimental profiling data.

Although these models all predict on-target activity and show moderate-to-high agreement, they were developed using different nucleases, guide designs, experimental readouts, training datasets, and machine learning frameworks. Consequently, CRISPR-HAWK reports their scores independently rather than combining them into a single consensus ranking. Detailed guidance on interpreting and selecting among these scores, together with a summary of each method’s nuclease system, PAM compatibility, training dataset, model architecture, and principal limitations, is provided in **Supplementary Data Section 2**.

#### 2.4.2 Predicting Residual On-Target Activity Across Variant Haplotypes

A critical challenge in variant-aware gRNA design is predicting how genetic variants at target sites affect the cutting efficiency of reference-designed guides. We address this by repurposing established off-target prediction models as surrogates for residual on-target activity when variants create mismatches. The key insight is that the same biophysical principles governing Cas9 binding and cleavage at mismatched off-target sites also apply when variants introduce mismatches at intended on-target sites.

CRISPR-HAWK implements this approach using two established models: cutting frequency determination (CFD) (Doench et al., 2016) and Elevation (Listgarten et al., 2018). While originally developed to predict off-target cleavage with sequence mismatches, these models quantify the fundamental relationship between sequence complementarity and Cas nuclease activity. When genetic variants alter a target site, they create imperfect base-pairing analogous to off-target mismatches.

For each reference-designed gRNA, CRISPR-HAWK identifies all alternative target sequences arising from genetic variants in the input population. These variant-containing sequences represent individual-specific on-target sites that differ from the reference. The tool then computes CFD and Elevation scores between each reference guide and its alternative targets. Lower scores indicate reduced predicted cleavage efficiency due to variant-induced mismatches, while scores near 1.0 suggest preserved activity despite genetic variation. This framework enables systematic assessment of how population-level genetic diversity may modulate therapeutic efficacy of CRISPR-Cas systems designed from reference genomes.

Among the available off-target scoring functions, CFD and Elevation were adopted for this purpose because both operate directly on guide-mismatched target pairs and expose an explicit, position- and type-dependent mismatch penalty that has been validated in variant-aware contexts (Cancellieri et al., 2023), making them directly applicable to residual on-target estimation. **Supplementary Data Section 3** provides a comparison with more recent deep learning-based off-target predictors, together with the rationale for this choice.

#### 2.4.3 Estimating Guide Specificity

To estimate guide specificity, CRISPR-HAWK integrates CRISPRitz (Cancellieri et al., 2020), a high-performance genome-wide search engine that identifies putative off-target sites allowing a user-defined number of mismatches and DNA/RNA bulges. For computational efficiency, off-target enumeration is performed against the reference genome, providing a baseline estimate of global specificity for each candidate guide. To extend this analysis to population-scale contexts, CRISPR-HAWK generates input files compatible with CRISPRme (Cancellieri et al., 2023), a variant- and haplotype-aware off-target nomination tool. This interoperability enables evaluation of guide specificity across diverse genetic backgrounds.

For each predicted off-target site, CRISPR-HAWK computes CFD and Elevation scores to estimate the likelihood of unintended cleavage. A global specificity score is then calculated for each gRNA by aggregating the CFDs of all its predicted off-targets, allowing users to rank and prioritize guides with minimal off-target potential.

#### 2.4.4 Annotating Guides

To support biological interpretation and downstream filtering, CRISPR-HAWK annotates candidate gRNAs with functional genomic context and sequence-derived features. Guides are annotated using user-provided genomic features, such as regulatory regions, coding exons, or disease-associated loci, like COSMIC cancer-related annotations (Sondka et al., 2024). Additionally, CRISPR-HAWK computes sequence-derived features that may influence gRNAs performance. GC content is calculated for each guide, as it has been shown to affect Cas binding affinity and gRNA stability (Yuen et al., 2017).

### 2.5 Reports Generation

CRISPR-HAWK generates a comprehensive set of output files and visualizations summarizing all candidate gRNAs and their associated features (**Figures 1H and 2**). The main output is a tabular report listing each candidate gRNA with its genomic coordinates, strand, PAM sequence, spacer sequence, predicted efficiency and specificity scores, functional and gene-level annotations, as well as haplotype and sample associations when variant data are provided (**Figure 2A**). When variant-aware analysis is performed, CRISPR-HAWK optionally produces a table reporting the reconstructed haplotypes for the input genomic region, including the constituent variants, population of origin, and allele frequencies (**Figure 2B**).

To facilitate data interpretation, the tool provides a series of graphical summaries (**Figure 2C**). These include comparative plots of on-target efficiency scores for reference-derived versus alternative guide sequences, illustrating the influence of genetic variation on predicted activity. Additional visualizations show the distribution of identified gRNAs by type (reference-only, variant-containing spacer, novel PAM, or both).

If candidate guide coordinates are supplied, CRISPR-HAWK generates a dedicated report restricted to the specified guides, detailing their corresponding alternative sequences arising from genetic variants, with associated residual on-target activity scores (**Figure 2D**). When the off-target search option is enabled, an off-target summary table is produced, listing all predicted off-target sites for each guide with their genomic positions, mismatch and bulge counts, and CFD/Elevation scores (**Figure 2E**). For each candidate guide, CRISPR-HAWK also produces a guide activity map that illustrates residual on-target activity across variant haplotypes with guide specificity, providing a global view of guide performance across diverse genomic backgrounds (**Figure 2F**).

## 3 Results

We evaluated CRISPR-HAWK across clinically relevant and well-characterized genomic regions to assess its ability to design and characterize variant- and haplotype-aware gRNAs using population-scale variant datasets. All analyses were performed using the GRCh38 human genome assembly, enriched with variants from three major population-scale datasets: the 1000 Genomes Project (1000G) (Consortium et al., 2015; Zheng-Bradley et al., 2017), the Human Genome Diversity Project (HGDP) (Bergström et al., 2020), and the Genome Aggregation Database (gnomAD) (Karczewski et al., 2020). The 1000G dataset includes whole-genome sequencing data from 2,504 individuals spanning 26 populations, organized into five superpopulations. The HGDP dataset comprises 929 individuals representing 54 populations, grouped into broader ancestries. The gnomAD dataset aggregates high-quality whole-genome sequencing data from 76,215 unrelated individuals, encompassing diverse genetic ancestries.

Using these datasets, we applied CRISPR-HAWK to two complementary tasks. First, we performed *de novo* guide enumeration and efficiency prediction at the BCL11A +58 erythroid enhancer, the target of sg1617, the clinically optimized guide used in Casgevy (exagamglogene autotemcel), one of the first FDA-approved CRISPR therapies for sickle cell disease and β-thalassemia. This analysis enabled both identification of alternative candidate guides and assessment of how population-level variants affect the predicted activity of sg1617 itself. Second, we quantified the effects of genetic variation on residual on-target activity and guide specificity across a broader panel of guides, including therapeutic gRNAs currently in clinical development and benchmark guides commonly used in off-target characterization studies.

To evaluate the computational scalability of CRISPR-HAWK, we benchmarked it evaluating the impact of key parameters on runtime and memory usage. Specifically, we measured performance across different genomic region sizes (1-100 kb), variant densities (0-20 variants/kb), phasing modes, and cohort sizes (1-2,000 samples). Overall, CRISPR-HAWK exhibited robust and consistent performance, maintaining scalability across all tested conditions without marked degradation. Comprehensive benchmarking results, including detailed runtime scaling with respect to each parameter and dataset size, are provided in **Supplementary Data Section 1**.

All analyses presented in this paper were performed on a Linux workstation with an AMD Ryzen Threadripper 3970X CPU with 32 cores and 64 GB of RAM.

### 3.1 Variant-Aware Guide Design on the BCL11A +58 Erythroid Enhancer

To demonstrate CRISPR-HAWK’s functionality, we focused on the BCL11A +58 erythroid enhancer, a regulatory region critical for hemoglobin gene expression (Bauer et al., 2013), targeted by the clinically optimized SpCas9 guide sg1617 (Frangoul et al., 2021; Canver et al., 2015; Wu et al., 2019). Analyzing the enhancer region (**Figure 3A**), CRISPR-HAWK identified several gRNA candidates on both the reference and alternative genomes (**Figure 3B**). Specifically, the tool found 295 gRNAs from 1000G and 268 from HGDP, with 40.3% and 34.3% containing genetic variants, respectively. A small subset (4.07% in 1000G; 2.61% in HGDP) originated from novel PAM motifs introduced by genetic variants and appeared exclusively in non-reference haplotypes. In contrast, the broader variant spectrum of gnomAD yielded a markedly higher fraction of guides carrying variants (94.28%), underscoring the importance of large-scale variant databases for comprehensive guide design (**Figure 3B**). Interestingly, using gnomAD variants CRISPR-HAWK identified four alternative gRNAs for sg1617.

**Fig. 3:**
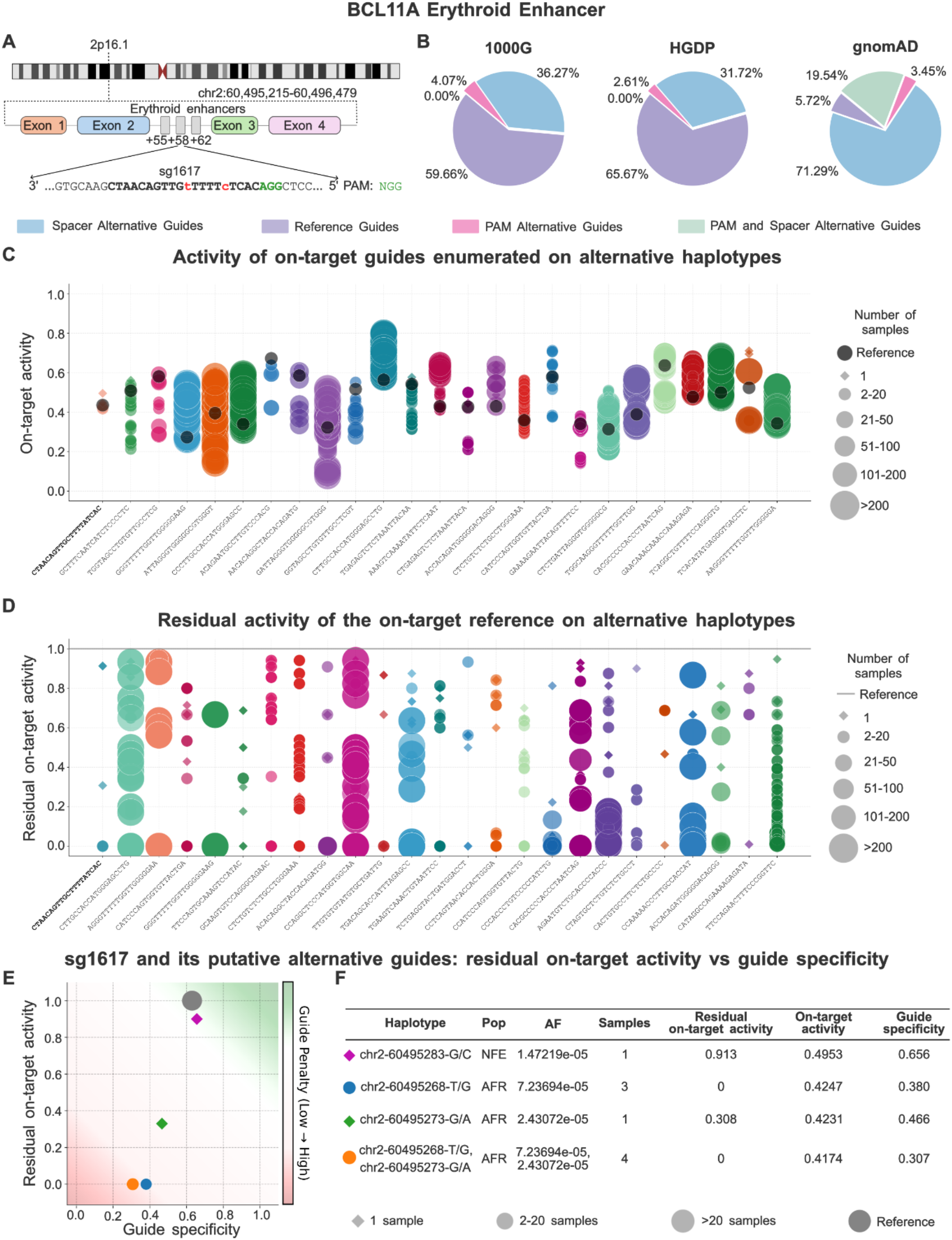
Variant-aware gRNA design and scoring analysis of the BCL11A +58 erythroid enhancer. **(A)** The clinically optimized guide sg1617 targets the +58 erythroid enhancer located within an intronic region of BCL11A. **(B)** Classification of gRNAs identified across three variant datasets (1000G, HGDP, and gnomAD): reference guides, spacer alternative guides (variants in spacer only), PAM alternative guides (variants creating novel PAM), and guides with variants in both PAM and spacer. **(C)** On-target efficiency scores (Azimuth) for the top 25 guides at the BCL11A enhancer, comparing reference sequences to haplotype-matched alternative sequences. Guides are ranked by maximum absolute score difference; sg1617 is shown in bold. Point size indicates the number of individuals carrying each variant. (D) Residual on-target activity (CFD) for the top 25 reference-designed guides at the BCL11A enhancer, showing predicted cleavage efficiency on variant-containing target sequences. Guides are ranked by maximum delta between reference and variant targets; sg1617 is shown in bold. Point size indicates the number of individuals carrying each variant. **(E)** Relationship between residual on-target activity and guide specificity for sg1617 and its variant-containing target sequences. Background shading indicates guide penalty, with green representing optimal combinations of high residual activity and high specificity. Off-target searches were performed with CRISPRme v2.1.7 using 1000G and HGDP variants, allowing up to 6 mismatches and 2 DNA/RNA bulges. **(F)** Summary of variant-containing target sequences for sg1617, reporting haplotype, population of origin (Pop), allele frequency (AF), sample count, residual on-target activity, on-target efficiency, and guide specificity.

We first assessed the on-target efficiency of guides designed to match specific haplotypes. By comparing the on-target efficiency scores predicted using the Azimuth model across the top 25 guides showing the largest score differences, we observed that specific variant combinations altered predicted activity, either enhancing or reducing efficiency depending on local sequence context (**Figure 3C**). Notably, a subset of alleles reduced predicted activity below 0.2, suggesting potential functional impairment. For sg1617 specifically, guides redesigned to perfectly match variant-containing sequences showed Azimuth scores comparable to the reference guide, indicating that variants in this region do not inherently compromise guide design potential. Although these predictions derive from computational models and may not fully reflect *in vivo* performance, they illustrate how genetic variants can modulate gRNA cutting efficiency. Dataset-specific efficiency estimates using Azimuth score for the BCL11A enhancer are provided in **Supplementary Data Section 4** and **Supplementary Figure 2**.

The preceding analysis assumes guides are redesigned to match each patient’s haplotype; however, in clinical practice, a single reference-designed guide is typically administered to all patients. To assess how variants affect the residual on-target activity of reference-designed guides, we computed CFD scores between each guide and its variant-containing target sequences (**Figure 3D**). In this context, variants function analogously to mismatches at off-target sites, potentially reducing or abolishing cleavage. Among the top 25 guides showing the largest CFD deltas, we observed a consistent reduction in predicted cleavage efficiency. Several alternative on-target sites displayed complete loss of activity (CFD = 0). Notably, this includes sg1617, the clinically deployed guide, which showed complete loss of predicted activity (CFD = 0) for two target sequences harboring gnomAD variants, suggesting that the approved therapy may have reduced or absent efficacy in patients carrying these variants.

Many of these variants occur at moderate to high population frequencies, implying that guides designed exclusively on the reference genome may fail to achieve effective editing in certain individuals. As with Azimuth, low CFD values do not imply total loss of cleavage but rather a reduced likelihood of efficient binding or cutting. Dataset-specific CFD results are detailed in **Supplementary Data Section 5** and **Supplementary Figure 3**.

To complete the characterization, we assessed the off-target potential of sg1617 and its alternatives using CRISPRme v2.1.7 (Cancellieri et al., 2023), integrating 1000G and HGDP variants and allowing up to six mismatches and two RNA/DNA bulges (Figures **3E**-**F**). Among the alternative target sequences (Figure 3F), those carrying the haplotypes chr2-60495268-T/G-chr2-60495273-G/A and chr2-60495276-G/A exhibited complete loss of predicted cleavage activity (CFD = 0), whereas the sequence carrying haplotype chr2-60495273-G/A retained moderate activity (CFD = 0.308). In contrast, the alternative target with haplotype chr2-60495283-G/C showed minimal impact on residual on-target activity (CFD = 0.913). Regarding guide specificity, most alternative sequences for sg1617 displayed a modest improvement in predicted specificity (**Figure 3E**). Notably, haplotype chr2-60495283-G/C maintained both high residual on-target activity (CFD = 0.913) and specificity comparable to the reference guide, whereas haplotypes with reduced on-target activity (CFD = 0) showed improved specificity, likely because the same mismatches that reduce on-target cleavage also reduce off-target binding.

To generalize these findings, we extended the analysis to two loci targeted by the CRISPR-Cas12a system, HBG1 and HBG2 (De Dreuzy et al., 2019). The analysis revealed similar patterns to those observed for BCL11A, where genetic variation modulated guide performance (**Supplementary Data Section 6** and **Supplementary Figures 4 and 5**).

Taken together, these results demonstrate that CRISPR-HAWK enables comprehensive assessment of guide performance across three dimensions: on-target efficiency for haplotype-matched designs, residual activity when reference guides encounter variant targets, and genome-wide specificity.

### 3.2 Evaluation of on-target activity of BCL11A +58 erythroid enhancer using multiple scoring models

Because CRISPR-HAWK reports on-target efficiency predictions from multiple models rather than a single consensus score (**Section 2.4.1**), we assessed the concordance among these independent estimates for haplotype-resolved guides. We scored all gRNAs identified at the BCL11A +58 erythroid enhancer using the five SpCas9 predictors supported by CRISPR-HAWK, namely Azimuth (Doench et al., 2016), RS3 (DeWeirdt et al., 2022), CRISPRon (Anthon et al., 2022), PLM-CRISPR (Hou et al., 2025), and sgDesigner (Hiranniramol et al., 2020), across sequences enriched with variants from the 1000G, HGDP, and gnomAD datasets (**Figure 4**). Predicted efficiencies varied across all predictors, indicating that sequence variation can markedly affect predicted editing activity irrespective of the scoring framework. Despite these differences in absolute predictions, the methods showed agreement in ranking high-performing candidates. Among the top-ranked guides, Azimuth (**Figure 4A**) shared five candidates with both RS3 (**Figure 4B**) and CRISPRon (**Figure 4C**), and six candidates with PLM-CRISPR (**Figure 4D**). sgDesigner (**Figure 4E**) showed the lowest overlap with the other predictors, with two to four shared guides, consistent with its distinct feature set and plasmid-based training data. Across all guides identified within the enhancer, pairwise Pearson correlation coefficients ranged from 0.58 to 0.87 (**Figure 4F**), indicating moderate-to-high concordance between methods. Higher correlations were generally observed between models with similar feature representations, whereas lower correlations reflected differences in model assumptions and training data. Overall, the moderate-to-high concordance between predictors supports the use of multiple scoring models in CRISPR-HAWK, which reports each prediction independently allowing users to select the most appropriate model.

**Fig. 4:**
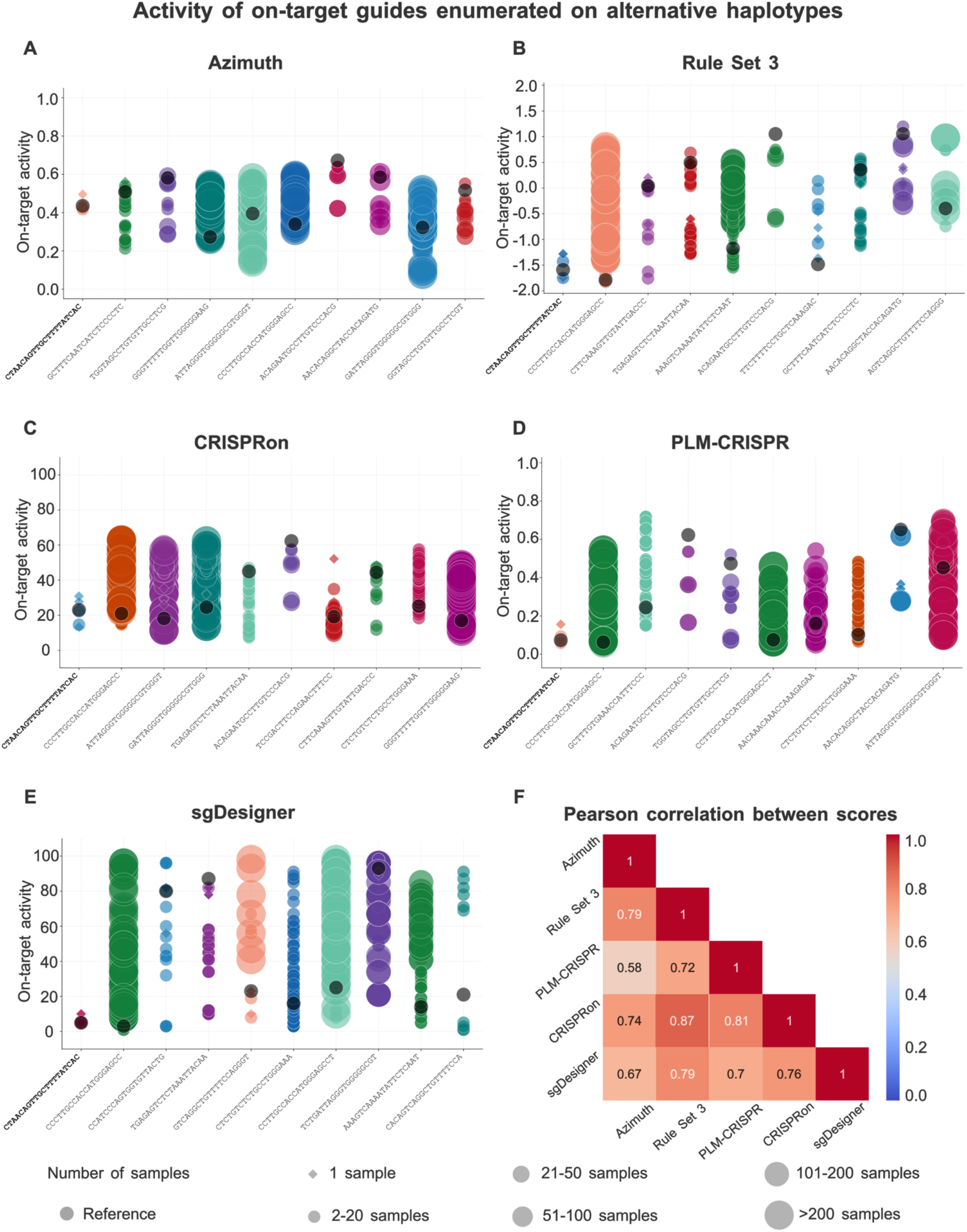
On-target activity scores are computed using Azimuth. **(A)**, Rule Set 3 **(B)**, CRISPRon **(C)**, PLM-CRISPR **(D)**, and sgDesigner **(E)**, using genetic variants from the 1000 Genomes Project, HGDP, and gnomAD datasets. Each point represents a guide RNA; point size reflects the number of samples carrying the corresponding variant(s), while diamonds denote singleton gRNAs (variant guides carried by single individuals). Reference guides are shown in dark, whereas haplotype-matched guides are color-coded. For each model, the top 10 guides ranked by the maximum absolute difference in predicted efficiency between reference and alternative haplotypes are displayed. **(F)** Heatmap of pairwise Pearson correlation coefficients between efficiency scores computed across all guides identified within the BCL11A enhancer region.

### 3.3 Residual On-Target Activity of Therapeutic and Benchmark gRNAs

We next extended the residual on-target activity analysis to additional genomic loci targeted by gRNAs in clinical or preclinical development (**Figure 5A**) including TRBC1/TRBC2 (Stadtmauer et al., 2020), HBB (DeWitt et al., 2016; Xu et al., 2019b), HBG1/HBG2 (Métais et al., 2019; De Dreuzy et al., 2019) and CCR5 (Xu et al., 2017; Xu et al. 2019a) genes. To provide broader context, we also included EMX1 and FANCF, benchmark loci commonly employed in off-target assessment studies (Tsai et al., 2015). For each guide, we computed CFD scores between the reference-designed sequence and all variant-containing target sequences identified across the three population datasets.

**Fig. 5:**
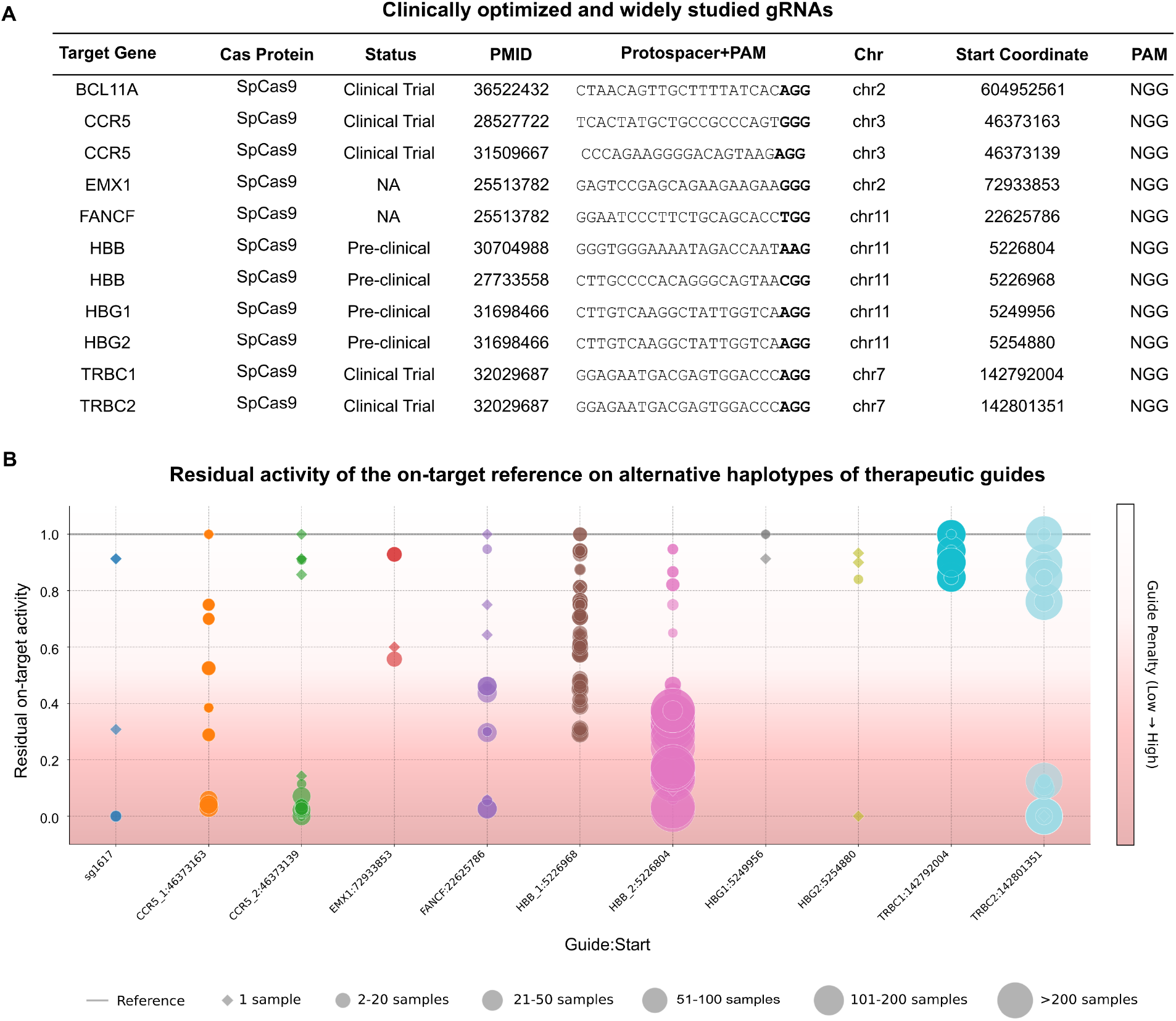
Residual on-target activity of therapeutic and benchmark gRNAs across variant-containing target sites. **(A)** Summary of analyzed guides, including target genes, Cas nucleases, clinical status, genomic coordinates, and PAM sequences. **(B)** CFD scores quantifying predicted cleavage efficiency of reference-designed guides on variant-containing target sequences. Each point represents a variant-containing target sequence; point size reflects the number of individuals carrying the corresponding variant(s).

For most analyzed guides, variants reduced predicted cleavage efficiency, with several cases showing a complete loss of activity (CFD = 0) at specific variant-containing target sites (**Figure 5B**). Notably, variants causing loss of cleavage activity often occurred in haplotypes shared by more than 200 individuals, as observed for HBB_2:5226804 and TRBC2:142801351 (**Figure 5B**). In contrast, other variants modulated gRNA activity more moderately, resulting in only partial decreases in CFD scores and suggesting a lesser impact on cleavage potential. For some guides, such as those targeting TRBC1, all variant-containing sites retained high predicted activity, indicating relatively low susceptibility to sequence variation at this locus.

Across the seven loci analyzed, 82.5% of reference-designed guides contained at least one variant predicted to modify on-target activity. Collectively, these results emphasize that even clinically validated gRNAs designed on the reference genome may experience substantial variability in predicted editing efficiency when applied to genetically diverse populations.

### 3.4 Validating CRISPR-HAWK sgRNA Predictions

To further validate CRISPR-HAWK on real genetic variation, we leveraged publicly available population-scale datasets to identify naturally occurring variants. As no public dataset currently combines on-target genome-editing measurements with genotype- or haplotype-resolved information for the same samples, direct experimental validation of on-target prediction is not feasible. For off-target predictions, we previously reported and experimentally validated variant-induced off-target effects for the BCL11A sg1617 guide in Cancellieri et al. (2023), and CRISPR-HAWK incorporates this validated framework via its integration with CRISPRme. To complement this with a model-independent assessment of on-target feasibility, we adopted a validation strategy grounded in a deterministic property of CRISPR-Cas systems, namely the strict requirement for an intact PAM. We assessed whether CRISPR-HAWK accurately identifies instances in which variants disrupt the canonical NGG PAM required by SpCas9, thereby abolishing targetability independently of any sequence-based predictive models. We identified multiple PAM-disrupting variants across diverse populations for which CRISPR-HAWK consistently predicts loss of valid targeting sites (see **Supplementary Data Section 7** for details). These cases identify individuals in the analyzed cohorts for whom current reference-based therapies are predicted to fail by deterministic argument alone, demonstrating that CRISPR-HAWK delivers clinically actionable findings even in the absence of variant-resolved on-target validation datasets.

## 4 Conclusions

CRISPR-HAWK’s core contribution consists of original engineering work specifically developed to enable variant- and haplotype-aware gRNA design at population scale. While CRISPR-HAWK leverages established off-target search engines (CRISPRitz, CRISPRme) and on-target scoring models, the methodological contributions at its core, namely efficient haplotype reconstruction from VCF data, haplotype-aware guide scanning with binary-encoded PAM matching, and per-nucleotide variant-to-spacer provenance tracking for residual on-target activity computation, are not available in any existing tool and cannot be reproduced through script-level chaining. Reconstructing per-individual haplotypes requires the sequential propagation of SNVs and indels along the target sequence while maintaining nucleotide-resolution coordinate mappings that remain consistent after each insertion or deletion event. In the absence of phasing information, heterozygous variants introduce combinatorial ambiguity that must be resolved efficiently. CRISPR-HAWK addresses this challenge through IUPAC-based encoding combined with selective enumeration restricted to candidate guide sequences, thereby avoiding the combinatorial explosion inherent to naïve approaches, and collapses biologically equivalent haplotypes via hashing-based deduplication to enable efficient processing of large cohorts. Computing residual on-target activity further requires a rigorous, per-nucleotide correspondence between each variant-containing spacer and its reference-derived counterpart, which cannot be reconstructed through post-hoc annotation of standard tool outputs. These components are tightly interdependent. Inaccuracies in indel offset propagation compromise downstream coordinate mappings. Incorrect haplotype deduplication increases computational redundancy, and loss of variant location undermines the interpretability of residual on-target activity scores. By encapsulating these interdependent processes within a unified and validated framework, CRISPR-HAWK enables reproducible and scalable population-aware gRNA design without requiring users to independently implement and validate each computational layer.

In this study, we introduced CRISPR-HAWK, a computational framework that integrates individual- and population-specific genetic variants and haplotype information into CRISPR-Cas gRNA design and evaluation. By accounting for genetic diversity, CRISPR-HAWK enables variant- and haplotype-aware gRNA design and assessment of three complementary performance metrics: on-target efficiency for haplotype-matched guides, residual on-target activity when reference-designed guides encounter variant targets, and genome-wide specificity, supporting the development of population-aware and personalized genome editing strategies.

CRISPR-HAWK’s core contribution consists of original engineering work specifically developed to enable variant- and haplotype-aware gRNA design at population scale. While CRISPR-HAWK leverages established off-target search engines (CRISPRitz, CRISPRme) and on-target scoring models, the methodological contributions at its core, namely efficient haplotype reconstruction from VCF data, haplotype-aware guide scanning with binary-encoded PAM matching, and per-nucleotide variant-to-spacer provenance tracking for residual on-target activity computation, are not available in any existing tool and cannot be reproduced through script-level chaining. Reconstructing per-individual haplotypes requires the sequential propagation of SNVs and indels along the target sequence while maintaining nucleotide-resolution coordinate mappings that remain consistent after each insertion or deletion event. In the absence of phasing information, heterozygous variants introduce combinatorial ambiguity that must be resolved efficiently. CRISPR-HAWK addresses this challenge through IUPAC-based encoding combined with selective enumeration restricted to candidate guide sequences, thereby avoiding the combinatorial explosion inherent to naïve approaches, and collapses biologically equivalent haplotypes via hashing-based deduplication to enable efficient processing of large cohorts. Computing residual on-target activity further requires a rigorous, per-nucleotide correspondence between each variant-containing spacer and its reference-derived counterpart, which cannot be reconstructed through post-hoc annotation of standard tool outputs. These components are tightly interdependent. Inaccuracies in indel offset propagation compromise downstream coordinate mappings. Incorrect haplotype deduplication increases computational redundancy, and loss of variant location undermines the interpretability of residual on-target activity scores. By encapsulating these interdependent processes within a unified and validated framework, CRISPR-HAWK enables reproducible and scalable population-aware gRNA design without requiring users to independently implement and validate each computational layer.

Applying CRISPR-HAWK to the BCL11A +58 erythroid enhancer, we demonstrated that naturally occurring variants can substantially alter gRNA sequence composition and predicted editing performance. Integration of population-scale variant data from 1000 Genomes, HGDP, and gnomAD revealed that a large fraction of candidate guides contains variants capable of modifying PAM recognition or spacer complementarity. Critically, analysis of sg1617, the guide used in Casgevy, revealed that certain variant-containing target sequences are predicted to completely abolish cleavage activity (CFD = 0), suggesting that the approved therapy may have reduced efficacy in individuals carrying these variants. Extending this analysis to therapeutic and benchmark gRNAs across seven loci confirmed these findings: 82.5% of reference-designed guides contained at least one variant predicted to modify on-target activity. These results emphasize the need for variant-aware evaluation in translational genome editing pipelines. Despite characterizing guide performance across these three axes, several limitations should be noted. CRISPR-HAWK relies on computational predictors that, while trained on experimental data, may not fully capture in vivo editing outcomes; experimental validation remains essential for clinical applications. The supported scoring models were developed and validated for specific Cas nucleases (Azimuth and CFD for SpCas9; DeepCpf1 for Cas12a); performance may vary when applied to other enzymes or engineered variants. The current implementation primarily addresses single nucleotide variants, with limited support for complex indels that may also affect guide performance. Additionally, predictions do not account for chromatin accessibility or local epigenetic context, which can substantially influence editing efficiency at specific loci. Delivery efficiency, which varies across cell types and delivery methods, is also beyond the scope of the current framework. The use of CFD and Elevation scores as surrogates for predicting residual on-target activity extends beyond their original scope. Both models were developed to quantify mismatch penalties at off-target genomic sites, and neither has been validated against variant-modified on-target sequences in genotype-resolved experimental or clinical settings. Their application here rests on the assumption that the same mismatch-tolerance mechanisms governing off-target cleavage also govern the reduction in on-target activity caused by sample-specific variants. Such estimates should therefore be interpreted as computational approximations of mismatch tolerance rather than predictors of therapeutic efficacy. Finally, population variant databases, while extensive, may not capture rare or population-specific variants relevant to individual patients.

From a translational perspective, the scoring models implemented in CRISPR-HAWK provide a practical framework for guide RNA selection. The first step is to assess targetability, as guides whose PAM sequence is disrupted by genetic variants are unlikely to support efficient editing and should therefore be excluded. Among the remaining candidates, predicted on-target efficiency, residual on-target activity in variant carriers, and variant-aware specificity provide complementary criteria for prioritization. Allele frequencies and population annotations further help assess the clinical relevance of residual variant sensitivity across different populations. Although the relative importance of these criteria depends on the intended application, prioritizing targetability before optimizing guide performance offers a rational and systematic strategy for both personalized and population-scale genome editing. A detailed description of this workflow is provided in **Supplementary Data Section 8**.

Overall, CRISPR-HAWK provides a comprehensive and scalable solution for assessing the functional consequences of genetic variation on CRISPR-Cas targeting. Its ability to incorporate haplotype and population-level variant data represents a methodological advancement toward precision genome editing, facilitating the development of guides optimized for diverse genetic backgrounds and enhancing the safety and efficacy of therapeutic interventions.

## 5. Data Availability

The CRISPR-HAWK source code (v0.1.2) is available at https://github.com/pinellolab/CRISPR-HAWK and https://github.com/InfOmics/CRISPR-HAWK. All data used to perform the analyses and generate the figures presented in this paper, including the CRISPR-HAWK guide reports and the off-target sites identified by CRISPRme for the sg1617 gRNA and its haplotype-matched alternatives, are available at https://doi.org/10.5281/zenodo.18070463. The scripts used to generate the results and figures reported in the manuscript are available at https://github.com/pinellolab/CRISPR-HAWK/tree/main/paper and at https://github.com/InfOmics/CRISPR-HAWK/tree/main/paper.

## 7. Author contributions statement

M.T., N.B., R.G., and L.P. conceptualized the methodology, M.T., A.K., G.C., and A.F developed the software. M.T., A.K., G.C., and A.F. analyzed the results, A.K., M.T., G.C., A.F., R.G., and L.P. wrote and reviewed the manuscript. R.G., and L.P. supervised the project.

## 8. Acknowledgments

The authors thank the members of InfOmics lab at University of Verona and of Pinello lab at Massachusetts General Hospital for their valuable suggestions.

## Supplementary information

Supplementary data are available at *Bioinformatics* online.

## Supplementary Data of

### 1 CRISPR-HAWK Computational Performance Assessment

To assess the scalability of CRISPR-HAWK and quantify the contribution of its binary-encoded PAM matching strategy to runtime and memory efficiency, we conducted a systematic benchmark covering a range of genomic region sizes, variant densities, phasing modes, and sample counts. All benchmarks were performed on a Linux workstation equipped with an AMD Ryzen ThreadRipper 3970X CPU (32 cores) and 64 GB of RAM.

Scoring steps (on-target efficiency and specificity/residual activity) were excluded from all benchmarks. These steps rely on external model APIs individually integrated within CRISPR-HAWK’s pipeline, represent the primary bottleneck of end-to-end runtime during typical analyses, and operate independently of the PAM matching strategy. Consequently, excluding them allows the benchmark to more accurately isolate and evaluate the computational performance of the core guide enumeration and variant-aware PAM matching components in CRISPR-HAWK.

Controlled benchmarking requires variant datasets with precisely defined density and phasing. We generated synthetic VCF files seeding variants on chromosome 2. Variant types were assigned stochastically, with single-nucleotide variants (SNVs) comprising 80% of variants, and insertions and deletions (1-5 bp) each representing 10%. To capture both typical and variant-dense genomic contexts, we systematically varied variant densities across five levels: 0 (reference only), 1, 5, 10, and 20 variants per kilobase. For phasing, separate VCFs were generated either with explicit haplotype phase information (phased) or with unresolved genotypes (unphased), the latter scenario inducing combinatorial haplotype expansion during gRNA sequence reconstruction. A fixed sample number of 100 was employed for benchmarks investigating the effects of region size and variant density.

CRISPR-HAWK encodes each haplotype using a 4-bit per-base representation supporting standard nucleotides and IUPAC ambiguity codes. This representation enables PAM detection through efficient bitwise AND operations rather than character-level string comparisons. Although both the binary-encoded implementation and a naïve character-based approach share the same asymptotic time complexity, *O*(*H* · *L*), with respect to the number of haplotypes *H* and the region length *L*, the binary representation substantially reduces constant factors in practice. In particular, the compact encoding reduces the memory footprint of haplotype storage by approximately 40-50% using a 4-bits-per-base representation of the encoded haplotype array.

To empirically quantify these advantages, we benchmarked the binary-encoded implementation against a naïve character-comparison approach across all combinations of region sizes and variant densities described above (**Supplementary Table 1** and **Supplementary Figure 1A—D**). Memory savings were consistent across all tested conditions. For example, at a region size of 100 kb with phased variants, binary encoding reduced peak memory by ∼42% at a variant density of 1 variant per kb (4.39 vs. 7.59 GB) and by ∼36% at 20 variants per kb (5.75 vs. 8.95 GB) (**Supplementary Figure 1A**). Runtime improvements were most pronounced at small-to-intermediate scales, reaching up to ∼1.7x at 10 kb and 1 variant per kb for phased data. At the highest tested variant density (100 kb / 20 variants per kb), runtimes of the two approaches converged (2,601 s vs. 2,743 s), reflecting a shift in the dominant computational cost from PAM matching to haplotype reconstruction (**Supplementary Figure 1B**). Notably, a density of 20 variants per kb substantially exceeds typical variant frequencies observed in the human genome (Consortium et al., 2015) and was included here primarily as a computational stress test.

Within parameter ranges most relevant for practical gRNA design, namely genomic regions of 1-20 kb with variant densities of 1-5 variants per kb, CRISPR-HAWK completes guide enumeration in under one minute using the binary-encoded implementation, with a peak memory footprint below 2GB. Comparable trends were observed when benchmarking unphased variant datasets, where binary encoding consistently reduced both memory consumption and runtime relative to naïve PAM-matching strategy (**Supplementary Figure 1C—D**). As in the phased case performance convergence between the two methods emerged only at the highest variant densities, where the combinatorial expansion of haplotypes required to reconstruct guides carrying unphased variants becomes the dominant contributor to runtime (**Supplementary Figure 1D**).

We next evaluated how runtime and memory scale with the number of samples in the input VCF. For this analysis, we considered a fixed genomic region of 10 kb with a variant density of 5 variants per kb and varied the number of samples from 1 to 2,000 under both phased and unphased configurations (**Supplementary Table 2** and **Supplementary Figure 1 E—F**).

Runtime increases approximately linearly with sample count. For phased inputs, scaling t from 1 to 1,000 samples resulted in a ∼350-fold increase in runtime (0.12 s to 41.59 s), consistent with the expected *O*(*H*) dependence on the number of reconstructed haplotypes (**Supplementary Figure 1F**). Analyses performed on unphased inputs were consistently faster than their phased counterparts at equivalent sample numbers (for example, 1,000 samples required 28.61 s compared to 41.59 s for phased data), reflecting the fact the unphased VCFs typically generate fewer distinct haplotypes per region during sequence reconstruction (**Supplementary Figure 1F**).

Memory consumption remained within practical limits across all tested conditions (**Supplementary Figure 1E**). Even at the largest scale evaluated, 2,000 phased samples for a 10 kb region, peak memory usage reached only 4.04 GB demonstrating that CRISPR-HAWK can efficiently process population-scale variant datasets while maintaining a modest memory footprint.

To complement the synthetic benchmarks, we profiled CRISPR-HAWK on a real genomic locus of clinical relevance: the BCL11A +58 erythroid enhancer, using variant datasets from the 1000 Genomes Project (1000G; 2,579 phased samples) (Consortium et al., 2015; Zheng-Bradley, 2019), the Human Genome Diversity Project (HGDP; 929 unphased samples) (Bergström et al., 2020), and the Genome Aggregation Database (gnomAD) (Karczewski et al., 2020) (**Supplementary Table 3** and **Supplementary Figure 1G—H**).

CRISPR-HAWK processed the 1000G and HGDP datasets in 3.66 s and 0.98 s, respectively, with memory usage remaining below 1 GB of memory in both cases. The analysis of the gnomAD dataset required 10.75 s despite involving fewer nominal samples (populations). This increased runtime is attributable to the substantial higher variant density in gnomAD, which leads to the reconstruction of a larger number of distinct haplotypes for PAM scanning.

Overall, these results demonstrate that CRISPR-HAWK maintains efficient runtime and modest memory requirements when applied to real population-scale datasets, confirming that variant- and haplotype-aware gRNA design can be performed on standard workstation hardware without the need for specialized high-performance computing resources.

**Supplementary Figure 1.**
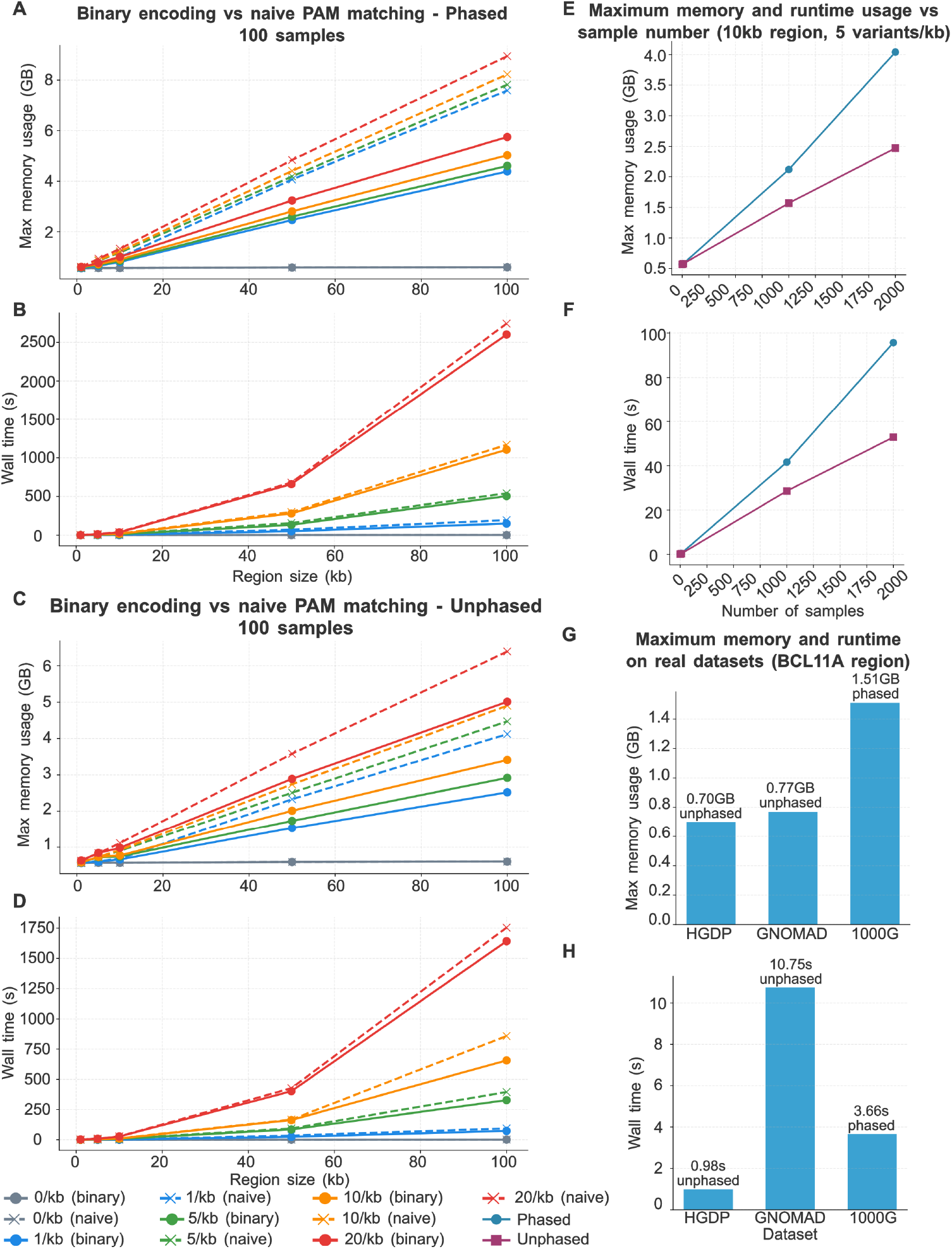
Performance benchmarking of CRISPR-HAWK across synthetic and real variant datasets. **(A)** Maximum memory usage comparison between the binary-encoded PAM matching strategy implemented in CRISPR-HAWK and a naïve character-comparison approach using phased variants (100 samples). **(B)** Wall clock runtime comparison between the binary-encoded PAM matching strategy implemented in CRISPR-HAWK and a naïve character-comparison approach using phased variants (100 samples). Benchmarks were performed across genomic sizes from 1—100 kb and variant densities ranging from 0 to 20 variants per kb. **(C)** Equivalent maximum memory usage benchmark using unphased variants. **(D)** Equivalent wall-clock runtime benchmark using unphased variants. **(E)** Peak memory usage scalability with respect to the number of samples in the input VCF using a fixed 10 kb genomic region with variant density of 5 variants per kb. **(F)** Wall-clock runtime scalability with respect to the number of samples in the input VCF using a fixed 10 kb genomic region with variant density of 5 variants per kb. **(G)** Maximum memory usage profiling on real population-scale datasets at the BCL11A +58 erythroid enhancer using variants from the 1000 Genomes Project, the Human Genome Diversity Project, and the Genome Aggregation Database. **(H)** Wall-clock runtime profiling on real population-scale datasets at the BCL11A +58 erythroid enhancer using variants from the 1000 Genomes Project, the Human Genome Diversity Project, and the Genome Aggregation Database.

**Supplementary Table 1.** Wall-clock time (seconds) and maximum memory usage (GB) for binary encoding and naïve PAM matching phased and unphased synthetic variants. The 0 variants per kb density represent reference-only runs (no VCF). Benchmarks were performed with 100 samples.

| Region | Variant-density | Phasing | Time-Binary (s) | Mem-Binary (GB) | Time-Naïve (s) | Mem-Naïve (GB) |
| --- | --- | --- | --- | --- | --- | --- |
| 1 kb | 0/kb | Reference | 0.02 | 0.56 | 0.03 | 0.56 |
| 1 kb | 1/kb | Phased | 0.14 | 0.57 | 0.14 | 0.57 |
| 1 kb | 1/kb | Unphased | 0.05 | 0.57 | 0.05 | 0.57 |
| 1 kb | 5/kb | Phased | 0.34 | 0.59 | 0.54 | 0.59 |
| 1 kb | 5/kb | Unphased | 0.19 | 0.58 | 0.31 | 0.58 |
| 1 kb | 10/kb | Phased | 1 | 0.61 | 1.56 | 0.61 |
| 1 kb | 10/kb | Unphased | 0.27 | 0.58 | 0.43 | 0.58 |
| 1 kb | 20/kb | Phased | 1.21 | 0.61 | 1.77 | 0.61 |
| 1 kb | 20/kb | Unphased | 2.29 | 0.63 | 2.33 | 0.62 |
| 5 kb | 0/kb | Reference | 0.07 | 0.57 | 0.08 | 0.57 |
| 5 kb | 1/kb | Phased | 1.91 | 0.66 | 3.26 | 0.72 |
| 5 kb | 1/kb | Unphased | 0.38 | 0.6 | 0.51 | 0.61 |
| 5 kb | 5/kb | Phased | 4.45 | 0.69 | 7.01 | 0.86 |
| 5 kb | 5/kb | Unphased | 3.31 | 0.72 | 4.64 | 0.72 |
| 5 kb | 10/kb | Phased | 8.9 | 0.72 | 8.4 | 0.87 |
| 5 kb | 10/kb | Unphased | 4.35 | 0.73 | 5.34 | 0.73 |
| 5 kb | 20/kb | Phased | 11.6 | 0.77 | 13.98 | 0.94 |
| 5 kb | 20/kb | Unphased | 9.28 | 0.84 | 10.03 | 0.84 |
| 10 kb | 0/kb | Reference | 0.13 | 0.57 | 0.25 | 0.57 |
| 10 kb | 1/kb | Phased | 4.82 | 0.81 | 8.19 | 1 |
| 10 kb | 1/kb | Unphased | 1.71 | 0.66 | 2.72 | 0.68 |
| 10 kb | 5/kb | Phased | 10.66 | 0.85 | 15.33 | 1.18 |
| 10 kb | 5/kb | Unphased | 7.23 | 0.74 | 9.52 | 0.9 |
| 10 kb | 10/kb | Phased | 16.21 | 0.91 | 20.74 | 1.23 |
| 10 kb | 10/kb | Unphased | 10.19 | 0.77 | 11.93 | 0.92 |
| 10 kb | 20/kb | Phased | 35.13 | 1.01 | 39.55 | 1.33 |
| 10 kb | 20/kb | Unphased | 26.78 | 0.98 | 27.11 | 1.11 |
| 50 kb | 0/kb | Reference | 0.67 | 0.59 | 0.91 | 0.59 |
| 50 kb | 1/kb | Phased | 50.5 | 2.47 | 70.68 | 4.07 |
| 50 kb | 1/kb | Unphased | 24.3 | 1.53 | 34.94 | 2.33 |
| 50 kb | 5/kb | Phased | 132.98 | 2.6 | 155.79 | 4.19 |
| 50 kb | 5/kb | Unphased | 84.99 | 0.98 | 93.04 | 2.51 |
| 50 kb | 10/kb | Phased | 279.67 | 2.81 | 298.68 | 4.41 |
| 50 kb | 10/kb | Unphased | 163.16 | 2.01 | 166.69 | 2.74 |
| 50 kb | 20/kb | Phased | 659.83 | 3.24 | 678.61 | 4.84 |
| 50 kb | 20/kb | Unphased | 401.44 | 2.89 | 426.28 | 3.58 |
| 100 kb | 0/kb | Reference | 1.18 | 0.6 | 1.53 | 0.6 |
| 100 kb | 1/kb | Phased | 149.69 | 4.39 | 193.11 | 7.59 |
| 100 kb | 1/kb | Unphased | 72.87 | 2.52 | 93.36 | 4.12 |
| 100 kb | 5/kb | Phased | 504.55 | 4.61 | 541.44 | 7.82 |
| 100 kb | 5/kb | Unphased | 327.74 | 2.92 | 394.63 | 4.47 |
| 100 kb | 10/kb | Phased | 1104.91 | 5.03 | 1167.48 | 8.23 |
| 100 kb | 10/kb | Unphased | 656.52 | 3.41 | 857.98 | 4.9 |
| 100 kb | 20/kb | Phased | 2601.01 | 5.75 | 2742.5 | 8.95 |
| 100 kb | 20/kb | Unphased | 1642.53 | 5.01 | 1755 | 6.39 |

**Supplementary Table 2.** Wall-clock time and peak memory as a function of sample count for phased and unphased inputs (10 kb region, 5 variants per kb density, binary encoding).

| Number of samples | Time – Phased (s) | Mem – Phased (GB) | Time – Unphased (s) | Mem – Unphased (GB) |
| --- | --- | --- | --- | --- |
| 1 | 0.12 | 0.57 | 0.13 | 0.57 |
| 10 | 0.5 | 0.58 | 0.36 | 0.58 |
| 100 | 10.66 | 0.85 | 7.23 | 0.74 |
| 1,000 | 41.59 | 2.12 | 28.61 | 1.57 |
| 2,000 | 95.74 | 4.04 | 52.84 | 2.47 |

**Supplementary Table 3.** Wall-clock time and peak memory for CRISPR-HAWK applied to the BCL11A +58 erythroid enhancer with three population-scale variant datasets, using binary-encoded PAM matching (SpCas9 NGG PAM, 20 nucleotides spacer). GnomAD is represented by 11 populations treated as samples. The higher runtime relative to sample number reflects the higher variant density of the gnomAD dataset.

| Dataset | Phasing | Number of samples | Time (s) | Mem (GB) |
| --- | --- | --- | --- | --- |
| 1000G | Phased | 2,579 | 3.66 | 0.7 |
| HGDP | Unphased | 929 | 0.98 | 0.77 |
| gnomAD | Unphased | 11* | 10.75 | 1.51 |

### 2 Guidance for the Interpretation of On-Target Activity Scores

CRISPR-HAWK reports several on-target activity scores for each candidate guide rather than single efficiency value. This choice reflects a basic property of the prediction task. In fact, On-target activity is an empirical quantity that no model measures directly and the available predictors were trained to approximate it under markedly different conditions. As summarized in **Supplementary Table 4**, the supported models differ in the nuclease and PAM for which they were developed, the length and composition of the guides used for training, the cell types and experimental readouts from which activity was derived, and the machine learning methodology used to relate sequence to activity. Each score therefore encodes a distinct set of biological and experimental assumptions and is best read as a complementary estimate of guide activity rather than as an interchangeable measurement of the same underlying quantity.

A direct consequence is that no single score is universally optimal. A model’s predictive accuracy depends on how closely the user’s nuclease, guide design, and cellular context resemble the data on which it was trained. Collapsing the individual predictors into a single consensus value would conceal the assumptions on which each score rests, assign comparable weight to models calibrated for unrelated experimental settings, and can yield rankings that appear authoritative yet are difficult to interpret or reproduce. For these reasons, CRISPR-HAWK does not compute a combined or consensus ranking. Instead, every applicable model is run independently and its score reported in parallel, leaving the selection and relative weighting of models to the user.

Selecting an appropriate score begins with the nuclease system, which constitutes a hard compatibility constraint rather than a preference. The scores are not transferable across enzymes. For example, DeepCpf1 (Kim et al., 2018) predicts activity for Cas12a guides and should be used only in that context, whereas the remaining models are intended for SpCas9 guides, with PLM-CRISPR (Hou et al., 2025) additionally accommodating several Cas9 orthologs. **Supplementary Table 4** specifies the nuclease and PAM compatibility of each model.

Within a compatible nuclease, the choice among models should be guided by the experimental context. Because the supported predictors were trained on different guide lengths and scaffolds, in different cell systems, and against different definitions of activity, the most informative score is generally the one whose training conditions most closely match the intended application. **Supplementary Table 4** reports the training dataset characteristics required to make this experimental design choice.

In practice agreement among several applicable models can be interpreted as added confidence in guide’s predicted activity, whereas a guide favored by only one model requires closer inspection of that model’s assumptions before prioritization. Such strong disagreement is, however, uncommon. Across guides identified in the BCL11A +58 erythroid enhancer, the SpCas9 models showed moderate-to-high pairwise agreement (Pearson R = 0.58—0.87), indicating that the choice of a particular score is rarely decisive for the top-ranked guides. This analysis is detailed in Section **3.2** of the main manuscript. Where scores do diverge, users are advised to favor the model whose training and experimental context best match their application rather than to average across models. Finally, all scores are *in silico* estimates. Therefore, a low predicted value reduces but does not categorically exclude measurable editing, and the scores are most appropriately used to prioritize candidates for experimental validation, rather than to make absolute accept-or-reject decisions.

**Supplementary Table 4.** Characteristics and interpretation of on-target activity prediction models integrated in CRISPR-HAWK. The table summarizes the six on-target scoring methods currently supported by CRISPR-HAWK, including their associated nuclease and PAM system, model architecture, training dataset characteristics, experimental setting, and year of publication. This information is intended to guide users in selecting and interpreting the most appropriate score for their nuclease system and experimental context.

| Method | Nuclease system | Model architecture | Training dataset description | Publication year |
| --- | --- | --- | --- | --- |
| Azimuth | SpCas9, NGG | GBRT | 2,500 sequences of 30nt; knockout/loss-of-function screen | 2016 |
| Rule Set 3 | SpCas9, NGG | GBRT | 46,526 sequences of 30nt + tracrRNA; knockout | 2022 |
| sgDesigner | SpCas9, NGG | GBRT+ SVM | 12,472 sequences of 53nt + scaffold; plasmid library/editing measured by sequencing | 2020 |
| CRISPRon | SpCas9, NGG | CNN | 23,902 sequences of 30nt; sgRNA activity measured with indels frequency | 2022 |
| PLM-CRISPR | 7 Cas9 variants, NGG | Protein-language model + CNN | 180,000 sequences of 59nt; sgRNA activity measured across Cas9 variants | 2025 |
| DeepCpf1 | Cas12a, TTTV | CNN | 1,251 sequences of 40nt; sgRNA activity measured with indels frequency + chromatin accessibility | 2018 |

**Supplementary Table 5.**
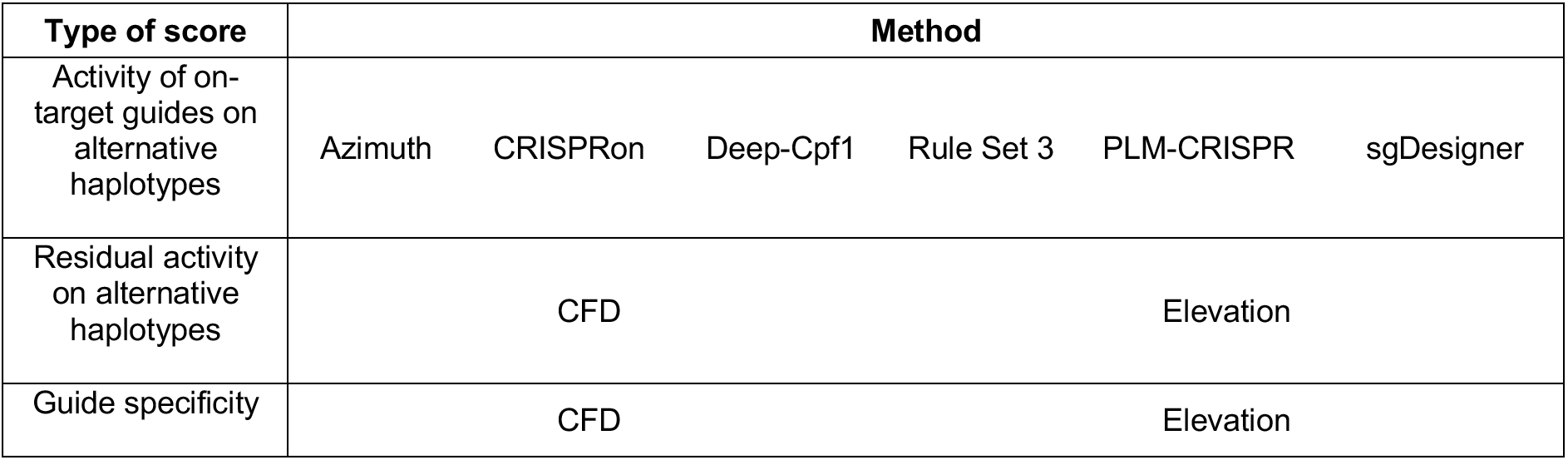
Scoring methods available in CRISPR-HAWK. Overview of the scoring functions implemented in CRISPR-HAWK, grouped by their functional role. On-target activity across alternative haplotypes is evaluated using multiple predictive models (Azimuth, CRISPRon, DeepCpf1, Rule Set 3, PLM-CRISPR, and sgDesigner). Residual activity on alternative haplotypes and guide specificity are assessed using CFD and Elevation scores, enabling comprehensive evaluation of both efficiency and specificity in a variant-aware context.

| Type of score | Method |  |  |  |  |  |
| --- | --- | --- | --- | --- | --- | --- |
| Activity of on-target guides on alternative haplotypes | Azimuth | CRISPRon | Deep-Cpf1 | Rule Set 3 | PLM-CRISPR | sgDesigner |
| Residual activity on alternative haplotypes | CFD |  |  | Elevation |  |  |
| Guide specificity | CFD |  |  | Elevation |  |  |

### 3 Repurposing Off-Target Activity Prediction Models to Evaluate Residual On-Target Activity

To quantify the impact of genetic variants on CRISPR-Cas on-target activity, we repurposed established off-target scoring frameworks to measure residual on-target activities. In this formulation, Cutting Frequency Determination (CFD) (Doench et al., 2016) and Elevation (Listgarten et al., 2018) scores are not used for their original purpose (off-target prioritization), but as proxies to estimate how genetic diversity within specific haplotypes or individuals alters the expected activity of an intended target site. This enables a variant-aware assessment of on-target efficiency by translating mismatch- or perturbation-dependent activity penalties into a quantitative reduction of canonical guide performance.

In principle, this formulation is not limited to CFD and Elevation. A broader class of off-target prediction models could be similarly repurposed to estimate variant-dependent perturbations in guide activity. This includes both classical and deep learning-based approaches (Zhang et al., 2023). Early neural architectures such as DeepCRISPR (Chuai et al., 2018) integrate convolutional encoders over guide-target sequences together with epigenetic features, enabling the joint modeling of sequence and chromatin determinants of cleavage activity. More recent models, including CRISPR-Net (Lin et al., 2020) and CRISPR-M (Sun et al., 2024), extend this framework by explicitly representing more complex guide-target interaction structures, such as DNA/RNA bulges and indel-associated alignments, and by leveraging recurrent or attention-based architectures to capture higher-order dependencies. Collectively, these methods demonstrate that off-target scoring functions can be generalized beyond mismatch-only representations to more expressive guide-genome interaction models.

Despite their conceptual applicability, these alternative approaches differ substantially in both formulation and interpretability. DeepCRISPR, for instance, learns a joint representation of sequence and epigenetic context using a convolutional neural network architecture. While it achieves high AUROC values, its improvements over CFD are relatively limited. Gains are more pronounced in AUPRC, indicating improved sensitivity in identifying true off-targets. However, this advantage is not consistently reflected in correlation-based metrics, suggesting limited improvement in capturing continuous variations in cleavage strength across sites.

CRISPR-Net extends DeepCRISPR’s paradigm by combining convolutional and recurrent components to better model structured dependencies in guide-target pairing, including bulge configurations. It achieves AUROC values comparable to CFD and Elevation, reflecting similar discriminative power between active and inactive sites, while showing improved AUPRC. CRISPR-M further advances this direction through an attention-based multi-view architectures that enables adaptive weighting of interaction features across mismatch and indel contexts. This results in improved performance over prior models, particularly in datasets enriched for indel-associated off-target events. When restricted to mismatch-only settings, CRISPR-M achieves performance comparable to CFD in AUROC and slightly improved AUPRC.

Among the available scoring strategies, CRISPR-HAWK uses CFD as the basis for residual on-target activity estimation. This choice is motivated by its experimental grounding and demonstrated ability to capture the functional consequences of sequence mismatches on Cas9 activity. Although originally developed for off-target prediction, CFD has been experimentally validated to accurately reflect the impact of genetic variation on cleavage efficiency, including in variant-aware contexts (Cancellieri et al., 2023). In addition, its explicit mismatch-position and mismatch-type penalty structure provides a transparent and biologically interpretable framework, which is particularly advantageous in large-scale variant-aware analyses where mechanistic interpretability is essential.

Elevation, in contrast, generalizes CFD by aggregating site-level off-target activities into a guide-level specificity score. Rather than modeling individual guide-target interactions in isolation, it integrates multiple CFD-derived estimates into a composite measure of global specificity. While this provides a useful abstraction for guide prioritization, it inherits the underlying assumptions of CFD-based activity modeling and does not fundamentally extend the representation of guide-target biophysics.

Finally, the design modular design of CRISPR-HAWK enables a seamless inclusion of heterogeneous off-target prediction models within a unified computational framework. This design choice ensures that CFD- and Elevation-based residual activity estimates can be naturally complemented or replaced by alternative classical or machine learning-based scoring functions as they mature.

### 4 Dataset-specific Evaluation of the On-Target Efficiency of BCL11A Enhancer Guides across Variant Haplotypes

By focusing CRISPR-HAWK’s analysis on the BCL11A +58 erythroid enhancer (Bauer et al., 2013), targeted by the clinically optimized guide sg1617 (Frangoul et al., 2021; Canver et al., 2015; Wu et al., 2019), we observed that genetic diversity significantly impacts predicted gRNA efficiency. We evaluated activity of on-target gRNAs retrieved across three variant datasets: 1000 Genomes (1000G) (Consortium et al., 2015), Human Genome Diversity Project (HGDP) (Bergström et al., 2020), and Genome Aggregation Database (gnomAD) (Karczewski et al., 2020). Each dataset was used independently to enrich the enhancer region with genetic variants (SNVs and short indels) with different levels of genetic diversity and sample representation.

For each dataset, we identified gRNA candidates on both the reference and alternative genomes. For each candidate guide we predicted its on-target activity using the Azimuth model (Doench et al., 2016). Notably, sg1617 displayed dataset-dependent alternative guide profiles. Enrichment with 1000G (**Supplementary Figure 2A**) and HGDP (**Supplementary Figure 2B**) variants produced no alternative gRNA for sg1617, indicating conservation of its target site in these cohorts. However, adding gnomAD variants (**Supplementary Figure 2C**) revealed four alternative gRNAs exhibiting comparable editing efficiencies, with Azimuth scores between 0.4 and 0.5. This reflects the broader genetic diversity captured by gnomAD, which includes substantially more individuals from a wide range of ancestries.

Beyond sg1617, we observed considerable variability in predicted efficiency among alternative guides. Some reference guides and their variant-derived forms showed similar Azimuth scores, whereas others displayed pronounced shifts in predicted activity depending on their variant content. These differences were consistently larger in the gnomAD dataset than in 1000G or HGDP. Although guides with low predicted efficiency occurred across all databases, the most substantial decreases, sometimes below 0.2, were primarily associated with alternative gRNAs carrying gnomAD variants. Such guides may exhibit reduced or negligible cleavage activity; however, these predictions are based solely on *in silico* modeling. A low Azimuth score does not categorically rule out measurable editing activity in experimental systems.

Overall, these results illustrate that gRNAs designed using a reference genome may show altered, and in some cases improved, performance when optimized for population- or sample-specific genomic contexts. Incorporating individual sequence variation directly into guide design can, under the right conditions, enhance predicted editing efficiency, underscoring the value of variant-aware strategies for personalized therapeutic applications.

**Supplementary Figure 2.**
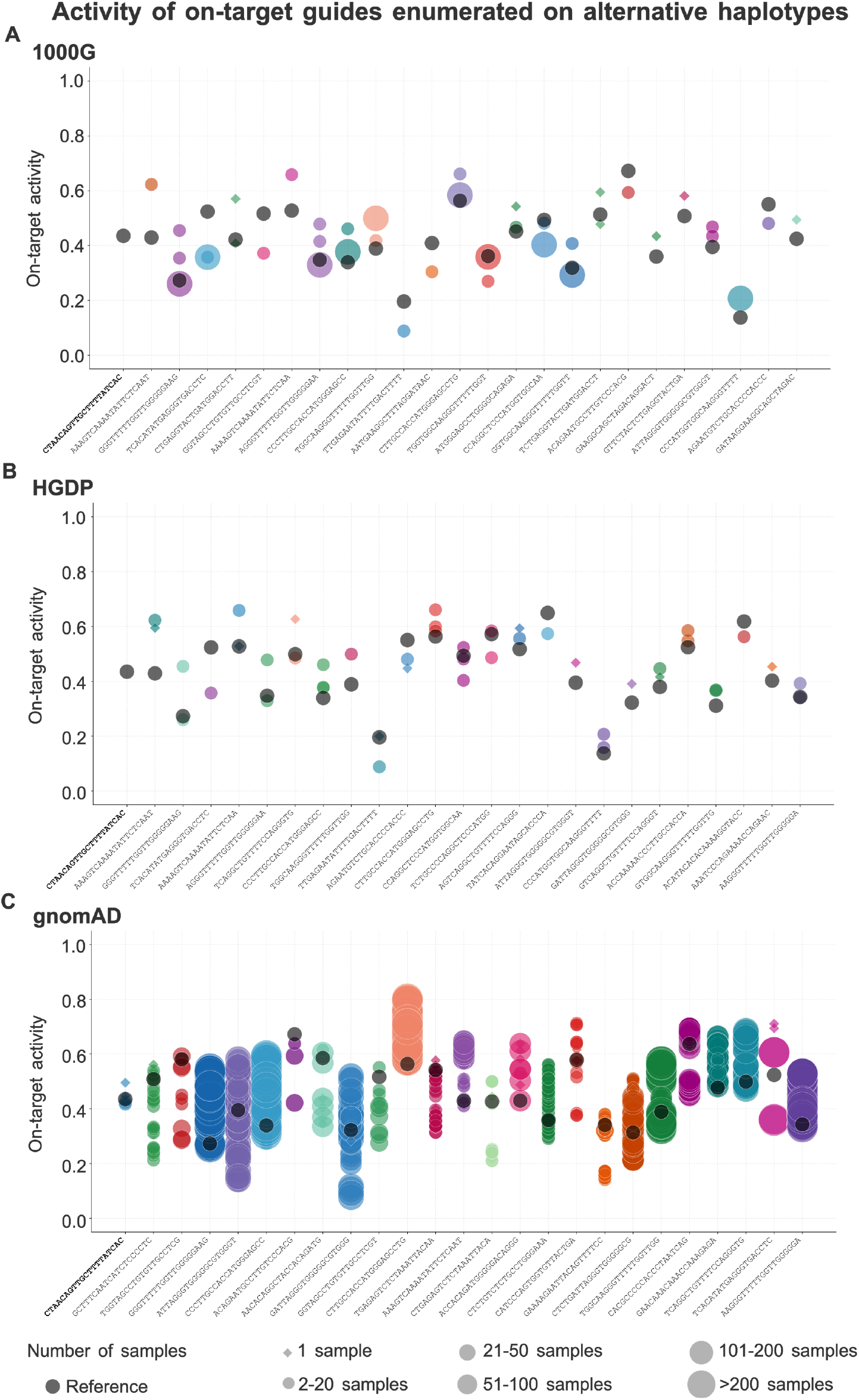
On-target efficiency of haplotype-matched guides across BCL11A enhancer. Analyses are based on variant data from 1000G (A), HGDP (B), and gnomAD (C). Each dot represents a guide, with dot size corresponding to the number of samples carrying the variant/s; diamonds indicate singletons. Grey dots represent reference guides, whereas colored dots represent variant-derived guides. The top 25 guides ranked by maximum absolute delta between reference and alternative guides are shown. On-target efficiency is evaluated using the Azimuth model.

### 5 Dataset-Specific Evaluation of Residual On-Target Activity across Variant Haplotypes

An important aspect in gRNA design is the extent to which guide performance deteriorates when applied to genetically diverse samples, particularly when variants fall within or near the target site. To assess this effect, we computed Cutting Frequency Determination (CFD) scores (Doench et al., 2016) between each reference guide and its corresponding alternative on-target sites within the BCL11A enhancer, independently enriched with variants from 1000G (**Supplementary Figure 3A**), HGDP (**Supplementary Figure 3B**), and gnomAD (**Supplementary Figure 3C**) datasets.

Across all datasets, we observed a general reduction in CFD scores for the top 25 guides exhibiting the largest discrepancies between reference and alternative targets. The magnitude of this reduction showed clear dataset-specific trends. When variant information from 1000G and HGDP was incorporated, most alternative targets maintained moderate CFD scores, typically above 0.4, indicating largely preserved binding and cleavage potential. In contrast, enrichment with gnomAD variants produced not only a higher number of alternative targets but also a substantially larger fraction with markedly reduced CFD scores. Many gnomAD-derived alternative on-target sites fell below 0.5, and several exhibited complete loss of predicted activity (CFD = 0), consistent with a potential abolition of on-target cleavage.

Of particular clinical relevance, two alternative on-target sites for the clinically optimized guide sg1617, identified exclusively using gnomAD variants, displayed CFD scores of zero, suggesting strong disruption of cleavage efficiency in those specific haplotypes. Conversely, another sg1617 alternative site showed no appreciable reduction in predicted activity, illustrating the heterogeneous impact of variation even within the same target region.

As with Azimuth predictions, CFD scores are generated by computational models; therefore, a predicted loss of activity does not definitively imply complete absence of cleavage in biological experiments. Nonetheless, these results highlight the importance of accounting for individual genetic variation. CRISPR-HAWK’s ability to systematically detect alternative target sites and quantify the predicted impact of variants provides users with a comprehensive and personalized assessment of guide performance across diverse genomic backgrounds.

**Supplementary Figure 3.**
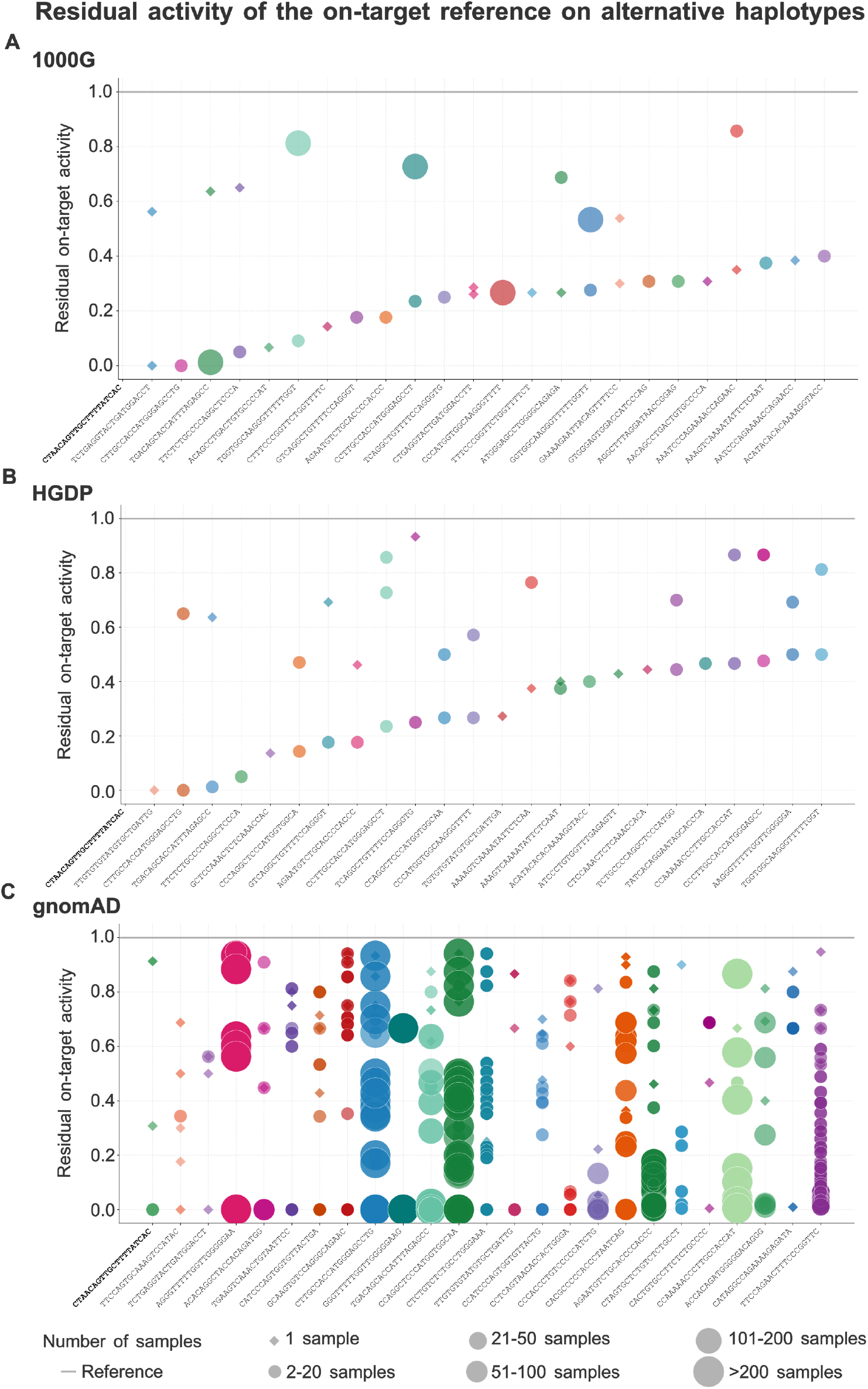
Residual on-target activity of haplotype-matched guides across BCL11A enhancer. Analyses are based on variant data from 1000G (A), HGDP (B), and gnomAD (C). Each dot represents a guide, with dot size corresponding to the number of samples carrying the variant; diamonds indicate singletons. The grey line represents reference guides, whereas colored dots represent variant-derived guides. The top 25 guides ranked by maximum delta between reference and alternative guides are shown. Variant effects on alternative on-targets are quantified using the CFD score.

### 6 Extraction of gRNAs and guide performance analysis for HBG1 and HBG2 genes targeted by CRISPR-Cas12a

To broaden our analysis beyond Cas9 systems, we applied variant-aware guide design and evaluation to the HBG1 (**Supplementary Figure 4**) and HBG2 (**Supplementary Figure 5**) loci (De Dreuzy et al., 2019), both targeted by CRISPR-Cas12a in therapeutic strategies aimed at inducing fetal hemoglobin expression. Retrieved gRNAs were assessed using variants from the 1000G, HGDP, and gnomAD datasets, which were integrated to enrich the DNA sequences of the target regions. To evaluate the contribution of different variant datasets to Cas12a guide design, gRNAs were categorized according to their variant content and database of origin (**Supplementary Figures 5A and 6A**).

For HBG1, CRISPR-HAWK identified 75 gRNAs from 1000G and 123 from HGDP, with 49.33% and 69.11% containing genetic variants, respectively. Notably, 2.66% of 1000G-derived guides and 8.94% of HGDP-derived guides originated from novel PAM motifs introduced by variants, representing sequences exclusive to non-reference haplotypes that would be missed in conventional reference-based design. For HBG2, 148 gRNAs were retrieved from 1000G and 241 from HGDP, with 45.95% and 66.80% harboring variants, respectively; 3.38% of 1000G and 7.88% of HGDP guides corresponded to variant-induced PAM motifs unique to alternative haplotypes. These proportions mirror those observed for HBG1, indicating similar patterns of variant impact across these closely related loci.

Analysis of the larger and more genetically heterogeneous gnomAD dataset revealed an even higher fraction of variant-containing gRNA candidates for both genes, with over 93% of retrieved gRNAs affected by genetic variation. Predicted on-target efficiencies were estimated using the DeepCpf1 model (Kim et al., 2018), a deep learning framework optimized for Cas12a activity, and guides were ranked according to the largest absolute difference in score between each reference guide and its variant-derived alternatives (**Supplementary Figures 5B and 6B**). The clinically optimized gRNA and its variant counterparts showed comparable predicted efficiencies, with DeepCpf1 scores ranging between 40 and 70 for both HBG1 and HBG2. Consistent with the BCL11A analysis, variant-derived guides exhibited both increases and decreases in predicted activity relative to reference-based designs, depending on the specific variant content. The magnitude and direction of these efficiency changes aligned with observations from Cas9-targeted regions.

Collectively, these results highlight the importance of incorporating population-specific variants in therapeutic gRNA design, as the majority of potential gRNAs deviate from the reference genome. The concordance of these findings across both HBG loci and across distinct CRISPR systems (Cas9 and Cas12a) reinforces the broad utility of variant-aware strategies in precision genome editing.

**Supplementary Figure 4.**
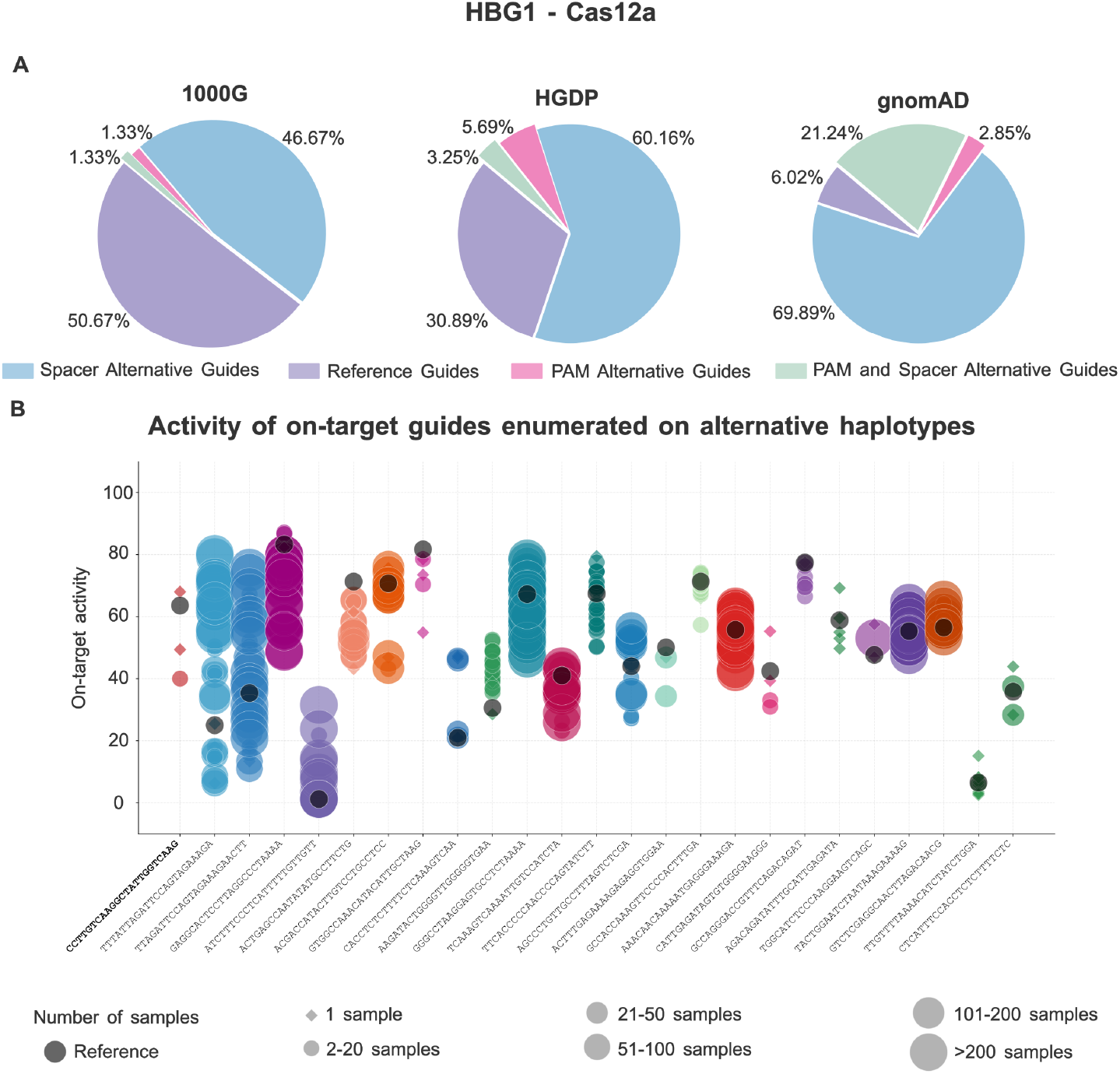
Variant-aware gRNA search and scoring analysis of a 2000 bp region of HBG1 gene centered on the therapeutic guide TTTGCCTTGTCAAGGCTATTGGTC. (A) Classification of retrieved guides across three variant datasets (1000G, HGDP, and gnomAD) showing dataset-specific proportions of reference and haplotype-matched guides, distinguishing those containing variants within the PAM, spacer, or both. (B) On-target efficiency of reference and haplotype-matched gRNAs across HBG1 evaluated using the DeepCpf1 model. Analyses are based on aggregated variant data from 1000G, HGDP, and gnomAD. Each dot represents a guide, with dot size corresponding to the number of samples carrying the variant/s; diamonds indicate singletons. Grey dots represent reference guides, whereas colored dots represent haplotype-matched guides. The top 25 guides ranked by maximum absolute delta between reference and alternative guides are shown.

**Supplementary Figure 5.**
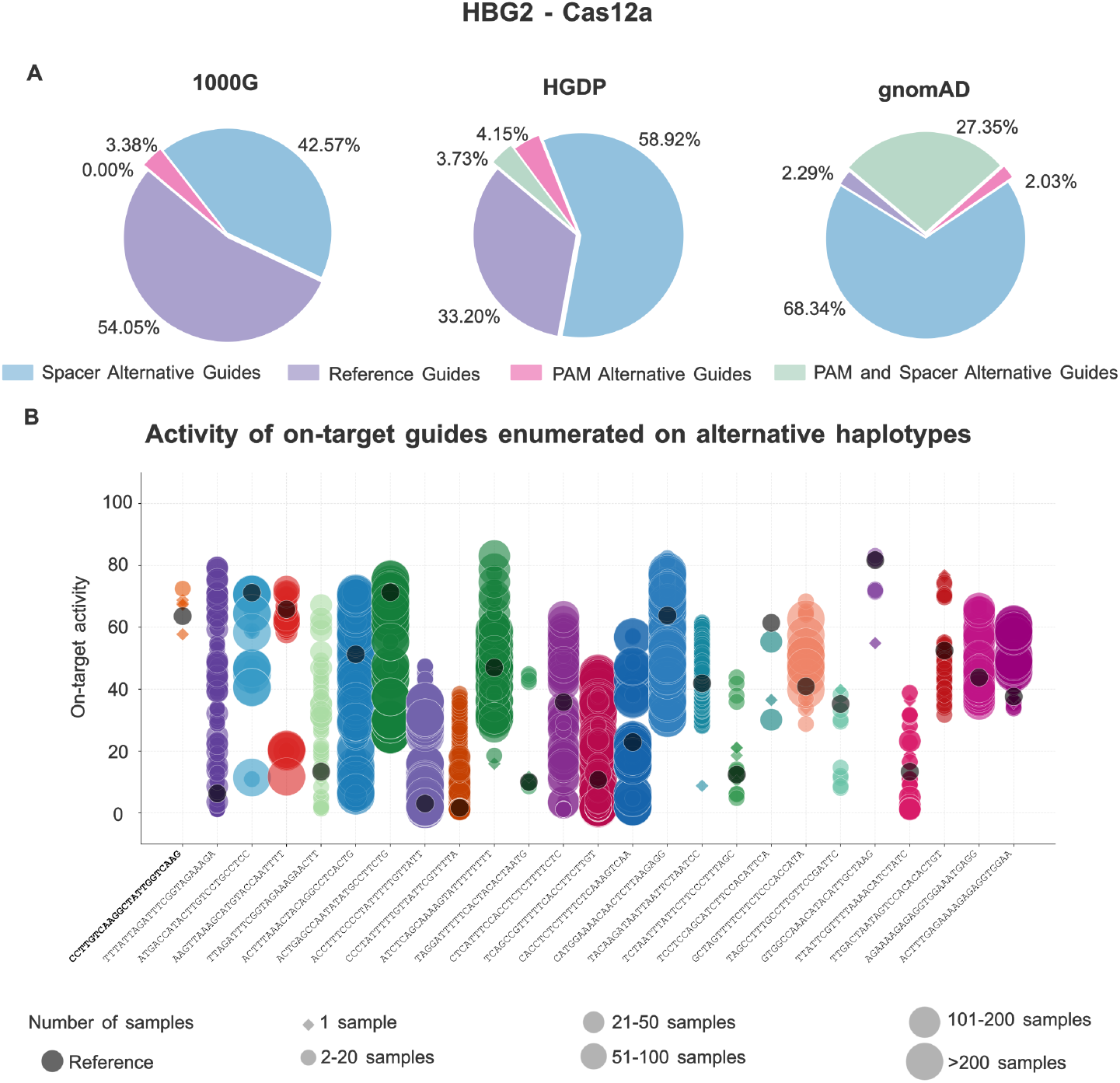
Variant-aware gRNA search and scoring analysis of a 2000 bp region of HBG2 gene centered on the therapeutic guide TTTGCCTTGTCAAGGCTATTGGTC. (A) Classification of retrieved guides across three variant datasets (1000G, HGDP, and gnomAD) showing dataset-specific proportions of reference and haplotype-matched guides, distinguishing those containing variants within the PAM, spacer, or both. (B) On-target efficiency of reference and haplotype-matched gRNAs across HBG2 evaluated using the DeepCpf1 model. Analyses are based on aggregated variant data from 1000G, HGDP, and gnomAD. Each dot represents a guide, with dot size corresponding to the number of samples carrying the variant/s; diamonds indicate singletons. Grey dots represent reference guides, whereas colored dots represent haplotype-matched guides. Top 25 guides ranked by maximum absolute delta between reference and alternative guides are shown.

### 7 Validating CRISPR-HAWK through variant-induced disruption of PAM sites on therapeutic targets

To further support the validity of CRISPR-HAWK predictions, we sought to evaluate its ability to capture variant effects on on-target activity using publicly available genome editing datasets. Ideally, such validation would rely on genome editing experiments performed on samples with known genotype or haplotype information. However, to the best of our knowledge, no publicly available datasets currently provide genome editing outcomes jointly with sufficiently resolved genetic variants at individual or haplotype level for the loci of interest.

In the absence of such data, we designed an alternative validation strategy based on a well-established mechanism and deterministic principle of CRISPR-Cas targeting: the strict requirement for an intact PAM. For SpCas9, disruption of the canonical NGG PAM abolishes nuclease binding and, consequently, editing activity, independently of any predictive model. This provides a robust validation scenario in which the expected outcome is unambiguous.

We therefore investigated whether CRISPR-HAWK can correctly identify cases in which genetic variants disrupt functional PAM sequences at therapeutically relevant and experimentally validated loci (see **Figure 5A** in main manuscript). Specifically, we searched for variants affecting the second or third nucleotide of the NGG motif, which are critical for PAM recognition.

To systematically identify such events, we applied CRISPR-HAWK using a two-step approach. First, we performed a relaxed gRNA search allowing fully degenerate PAM (NNN), thereby identifying all candidate protospacers regardless of PAM compatibility. Second, we compared these results with those obtained under the canonical NGG constraint within the same genomic regions. This comparison enabled the detection of instances in which sequence variation converts a functional NGG PAM into a non-functional motif.

Using this strategy, we identified multiple PAM-disrupting variants for gRNAs targeting CCR5 and HBB supported by population-scale datasets, including gnomAD and the Human Genome Diversity Project. These variants occur in real individuals across diverse populations and, in some cases, are observed in haplotype-resolved samples. For individuals carrying such variants, CRISPR-HAWK correctly predicts the absence of valid targeting sites due to the loss of a functional PAM, implying a complete failure of SpCas9-mediated editing at those loci.

The full list of disrupted PAMs, including genomic coordinates, sgRNA sequences, strand information, GC content, associated variants, allele frequencies, and population annotations, is reported in **Supplementary Table 6**.

This analysis provides a concrete validation of CRISPR-HAWK’s ability to capture the functional consequences of genetic variation on genome editing outcomes. Importantly, it demonstrates that the framework accurately identifies variant-induced loss of targetability in real genomic data, even in the absence of direct experimental editing measurements. These results reinforce the importance of incorporating genotype- and haplotype-aware analyses for the reliable design of CRISPR-based therapeutic strategies.

**Supplementary Table 6.**
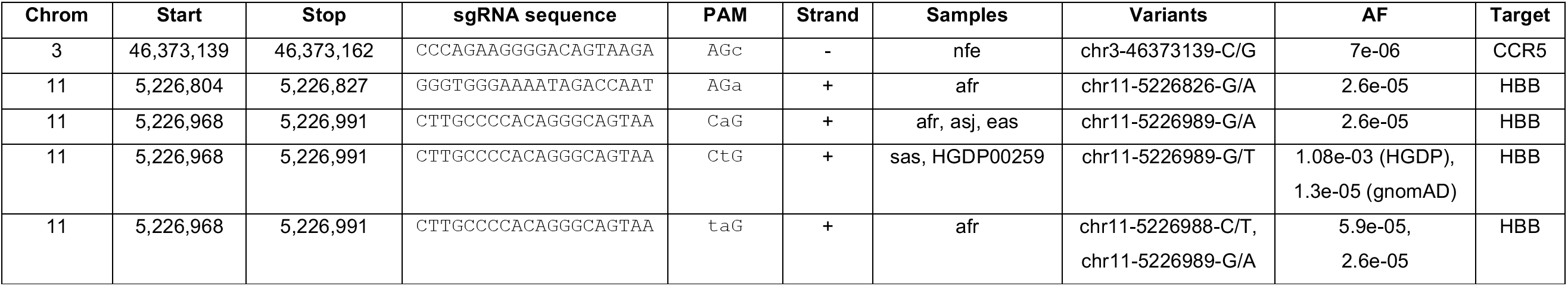
Variant-induced disruption of PAM sequences in therapeutically relevant guides targeting CCR5 and HBB. Genetic variants affecting the second or third nucleotide of the canonical NGG PAM are reported. These substitutions convert functional PAMs into non-canonical motifs, thereby abolishing SpCas9 binding and cleavage. For each candidate guide, we report genomic coordinates, sgRNA sequence, disrupted PAM, strand orientation, populations/samples in which the variant is observed, variant identifier, allele frequency (AF), and associated target. Allele frequencies are derived from gnomAD unless otherwise specified. HGDP frequencies are reported when available.

### 8 Recommended Workflow for Prioritizing Candidate Guides

CRISPR-HAWK reports multiple complementary metrics describing distinct aspects of guide performance, including targetability (PAM integrity in the presence of genetic variants), predicted on-target efficiency, residual on-target activity on variant-containing target sequences (estimated using CFD (Doench et al., 2016) and Elevation (Listgarten et al., 2018)), variant-aware off-target specificity, and the population frequency and ancestry distribution of variants affecting guide performance. Because these metrics capture different biological properties rather than alternative estimates of the same phenomenon, they should not be collapsed into a single composite score. For example, a guide predicted to be highly efficient cannot compensate for the complete loss of targetability caused by disruption of its PAM sequence. Accordingly, CRISPR-HAWK reports these metrics independently and leaves their integration to the specific biological or clinical application.

For guide prioritization, we recommend beginning by defining the genetic context relevant to the intended application, ranging from an individual genotype for personalized genome editing to population-scale datasets such as 1000 Genomes (Consortium et al., 2015), HGDP (Bergström et al., 2020), or gnomAD (Karczewski et al., 2020) for population-aware guide design. Haplotype reconstruction should precede guide evaluation so that all subsequent analyses are performed on the genetic backgrounds of interest. Candidate guides whose canonical PAM is disrupted by genetic variants should then be excluded, as the loss of a functional PAM abolishes nuclease recognition irrespective of any predicted efficiency score. This situation is exemplified by our analyses of the CCR5 and HBB loci, where common variants convert canonical PAMs into non-functional motifs (**Supplementary Data Section 7** and **Supplementary Table 6**).

Among the remaining targetable guides, priority should be given to candidates predicted to maintain high on-target activity across the reconstructed haplotypes using an appropriate nuclease-specific model. Guides predicted to perform well only on the reference genome should be deprioritized because their activity may not generalize across genetically diverse individuals. Residual on-target activity should then be evaluated by comparing each reference guide with its corresponding variant-containing target sequences using CFD or Elevation. Guides exhibiting substantial reductions in predicted activity across common haplotypes are less suitable for broad application, as illustrated by the therapeutic BCL11A sg1617 guide, for which certain gnomAD-derived haplotypes markedly reduce predicted activity whereas others have little or no effect.

Variant-aware specificity should subsequently be considered, particularly in therapeutic settings where minimizing unintended editing remains a key safety objective. Finally, allele frequencies and population annotations provide an essential framework for interpreting the potential clinical impact of variant-associated changes in guide performance, allowing users to prioritize guides that maintain robust activity across common variants and diverse ancestral groups.

Overall, this workflow identifies guides that remain targetable across the genetic backgrounds of interest, maintain high predicted on-target activity, preserve activity in the presence of common genetic variants, and exhibit acceptable variant-aware specificity. We emphasize that this workflow is intended as a recommended strategy rather than a prescriptive decision framework. The relative importance of on-target efficiency, residual activity, specificity, and population context may be adjusted according to the biological question and intended application, whereas targetability should always be treated as a mandatory prerequisite. Importantly, all the required metrics are computed directly by CRISPR-HAWK, allowing this workflow to be applied without any additional analyses beyond a standard software run.

